# Integrated solid/solution NMR assignment allows mapping dynamics and ligand binding in a 134 kDa enzyme

**DOI:** 10.64898/2026.07.22.740015

**Authors:** Federico Napoli, Rajkumar Singh, Anna Kapitonova, Virgil Aitenbichler, Giorgia Toscano, Barbara Perrone, Paul Schanda

## Abstract

Understanding enzyme function requires characterizing not only static structure but also dynamics and ligand interactions. NMR spectroscopy provides this insight at atomic resolution, yet for large proteins the difficulty of resonance assignment has largely confined such studies to systems below *∼*50 kDa, or to observing only methyl groups. Here we present an integrated magic-angle spinning (MAS) and solution NMR study of the 134 kDa tetrameric malate dehydrogenase from *Ignicoccus islandicus* (*Ii*MDH), an enzyme of particular interest as an evolutionary intermediate between allosteric lactate dehydrogenases and non-allosteric malate dehydrogenases. By combining high-dimensional (up to 4D) MAS NMR experiments on sedimented protein with solution NMR, we achieved 92% backbone heavy-atom assignment and 91% assignment of all Ile-δ_1_, Leu-δ_1_/-δ_2_, Val-γ_1_/-γ_2_, Met-ɛ and Thr-γ methyl groups. Building on these assignments, MAS NMR ^15^N rotating-frame relaxation (*R*_1_*_p_*) measurements revealed pronounced microsecond-timescale backbone dynamics in functionally critical regions, including the catalytic loop and the mobile surface loop. Complementary methyl-axis order parameters from solution NMR identified additional flexible sites in the hydrophobic core. Chemical shift perturbation experiments upon addition of the substrate analogue oxamate, monitored via backbone ^1^H-^15^N TROSY, revealed both active-site contacts and rearrangements of helices a2F and a3G, regions implicated in allosteric signal transmission. The integrated approach demonstrated here exploits the distinct strengths of MAS and solution NMR, and provides a comprehensive view of structure, dynamics, and substrate interactions in a large oligomeric enzyme that would not be accessible by either technique alone.

## Introduction

Enzyme catalysis depends not only on a single “static” three-dimensional structure, but critically relies on the enzyme’s ability to sample alternate conformations that are transiently populated during the catalytic cycle or upon ligand binding^1,2^. This is particularly evident in allosteric enzymes, where binding of a substrate or effector at one site modulates activity at a distant site through structural and dynamical changes that propagate across the protein. Understanding the mechanistic origins of allostery therefore requires characterizing not only the ground-state structure – where X-ray crystallography and cryo-EM excel –, but also the dynamics and interactions of the protein at atomic resolution^3,4^.

The superfamily of malate and lactate dehydrogenases (MDHs and LDHs) offers a compelling system for studying the origins of allostery^5,6^. LDHs and MDHs share the same fold and catalyze chemically equivalent reactions, yet LDHs exhibit allosteric regulation between their four subunits while MDHs do not^7^. The malate dehydrogenase from the archaeon *Ignicoccus islandicus* (*Ii* MDH) occupies a particularly interesting position in the phylogeny of this superfamily, sitting at the evolutionary intersection between allosteric LDHs and non-allosteric MDHs^7^. As a 134 kDa homotetramer, *Ii* MDH catalyzes the reduction of oxaloacetate to malate coupled to the oxidation of NADH to NAD^+^, within a conserved active site that brings cofactor and substrate into close proximity. Interestingly, it also catalyses the conversion of pyruvate to lactate – the reaction performed by LDHs – with signs of allosteric regulation, albeit at lower efficiency^8^. These properties make *Ii* MDH an ideal model to dissect the structural and dynamical determinants of allostery at the atomic level.

Nuclear magnetic resonance (NMR) spectroscopy is uniquely suited to this endeavour: it reports on both time-averaged structure and internal dynamics at the resolution of individual atoms, and it can probe proteins directly in solution, where transient interactions with substrates and cofactors are accessible^9,10^. However, solution NMR has long faced an inherent size limitation, because slow Brownian tumbling leads to rapid signal loss during the coherence-transfer steps of multidimensional experiments. TROSY-based backbone^11^ ^1^H-^15^N and methyl^12^ ^1^H-^13^C correlation experiments have substantially alleviated this problem. Methyl-TROSY approaches in particular have provided key functional insights into molecular machines approaching the megadalton range^13,14^. Nonetheless, important practical limitations remain. First, methyl-TROSY experiments report exclusively on methyl-bearing side chains, leaving polar residues — which are often directly involved in active sites and allosteric interfaces — invisible. Second, and more fundamentally, while 2D ^1^H-^15^N TROSY spectra can be obtained for large proteins, *de novo* resonance assignment requires usually 3D spectra with ^15^N-^13^C and ^13^C-^13^C coherence-transfer steps, but such experiments become prohibitively inefficient at high molecular weight. Common solutions to this problem fall into three categories^15^: (i) in some cases, large proteins may be dissected into smaller domains, which can be assigned, and the assignment may subsequently be transferred to the spectra of the entire protein^16–20^. The process is bound to the ability to track resonances from the small subdomains to the entire complex – despite the necessarily broken interfaces –, and often requires extensive trial-and-error testing of constructs. (ii) As through-space polarization transfer (NOE) is efficient for large proteins, NOESY spectra along with the known 3D structure may also allow for resonance assignment; this technique is often used for methyl assignments^21–25^. (iii) Alternatively, mutagenesis-based assignment uses a series of 2D ^1^H-^13^C spectra of single-point mutants whereby the replacement of a methyl-bearing residue leads to the disappearance of a single cross peak^20,26,27^. Besides being laborious, mutations change the protein, which can lead to additional unwanted effects, such as peak shifts of non-mutated residues.

Magic-angle spinning solid-state NMR (MAS NMR) offers a complementary and in several respects advantageous approach. By replacing Brownian tumbling with mechanical sample rotation, MAS NMR removes the molecular-weight dependence of resolution and sensitivity, and proteins of hundreds of kilodaltons — frequently in oligomeric assemblies — have been studied by MAS NMR to reveal their structures, dynamics and interactions^28–30^. Large protein assemblies can be converted into a suitable “sedimented” solid-state sample by ultracentrifugation^31–33^, while remaining highly hydrated (typically more than 50% water by volume) and retaining structure and dynamics closely similar to those in solution. This makes sedimented samples well suited for backbone resonance assignment and for measurements of site-specific dynamics across a wide range of timescales^34,35^. However, the study of transient intermolecular interactions — such as substrate binding to an enzyme — is less straightforward in sedimented samples.

Here we exploit the complementary strengths of MAS and solution NMR to obtain a comprehensive view of the structure, dynamics and substrate interactions of *Ii* MDH. We show that high-dimensional MAS NMR experiments on the sedimented protein yield a near-complete backbone resonance assignment, which is then transferred to the methyl groups through a series of solution– and MAS-NMR experiments, allowing for the unambiguous assignment of 91.3% of Ile-δ_1_, Leu-δ_1_/-δ_2_, Val-γ_1_/-γ_2_, Met-ε and Thr-γ methyl groups. The combined assignment enables several parallel lines of investigation of dynamics: ^15^N rotating-frame relaxation (*R*_1_*_p_*) measurements by MAS NMR reveal microsecond-timescale backbone dynamics in functionally critical regions, including the catalytic loop and a mobile surface loop which is involved in the coordination of substrates; methyl-directed order parameter measurements and rotamer-equilibrium assessments in solution provide additional insights into side chain mobility on the ps-ns time scale. Backbone ^1^H-^15^N TROSY spectra in solution – assigned through the MAS-NMR data – map the structural response to the substrate analogue oxamate across the structure, hinting to inter-subunit allosteric communication. Together, these results demonstrate that the integrated MAS/solution NMR strategy presented here is a powerful and practical route to studying dynamics and interactions in large oligomeric proteins that are inaccessible to either technique alone.

## Results and discussion

### High-resolution MAS and solution NMR of the *Ii* MDH enzyme

Figure 1a-c compares TROSY-based solution-state two-dimensional (2D) ^1^H-^15^N and 3D HNCO and HNCA spectra of deuterated (U-^2^H,^13^C,^15^N) *Ii* MDH with equivalent spectra recorded by MAS NMR techniques on a sedimented protein. The first important observation from this comparison is that the cross-peak positions are very similar in solution and in the sediment, mirroring previous reports^32,36^. This establishes that data from both approaches can be combined and complement each other. Crucially, the number of detectable cross-peaks in solution is significantly lower than in MAS, particularly in the 3D experiments, in which the signal is lost during the ^15^N-^13^C coherence-transfer delays. This signal loss is even more pronounced in experiments involving additional ^13^C-^13^C transfer steps, such as HNcoCACB or HNCACB, such that these crucial experiments for the assignment are practically useless for the *Ii* MDH case (data not shown). In contrast, in the sedimented sample, excellent spectral quality was found in numerous high-dimensional experiments, establishing MAS-NMR as the way forward for the assignment of the backbone and the side-chain heavy atoms.

**Figure 1:**
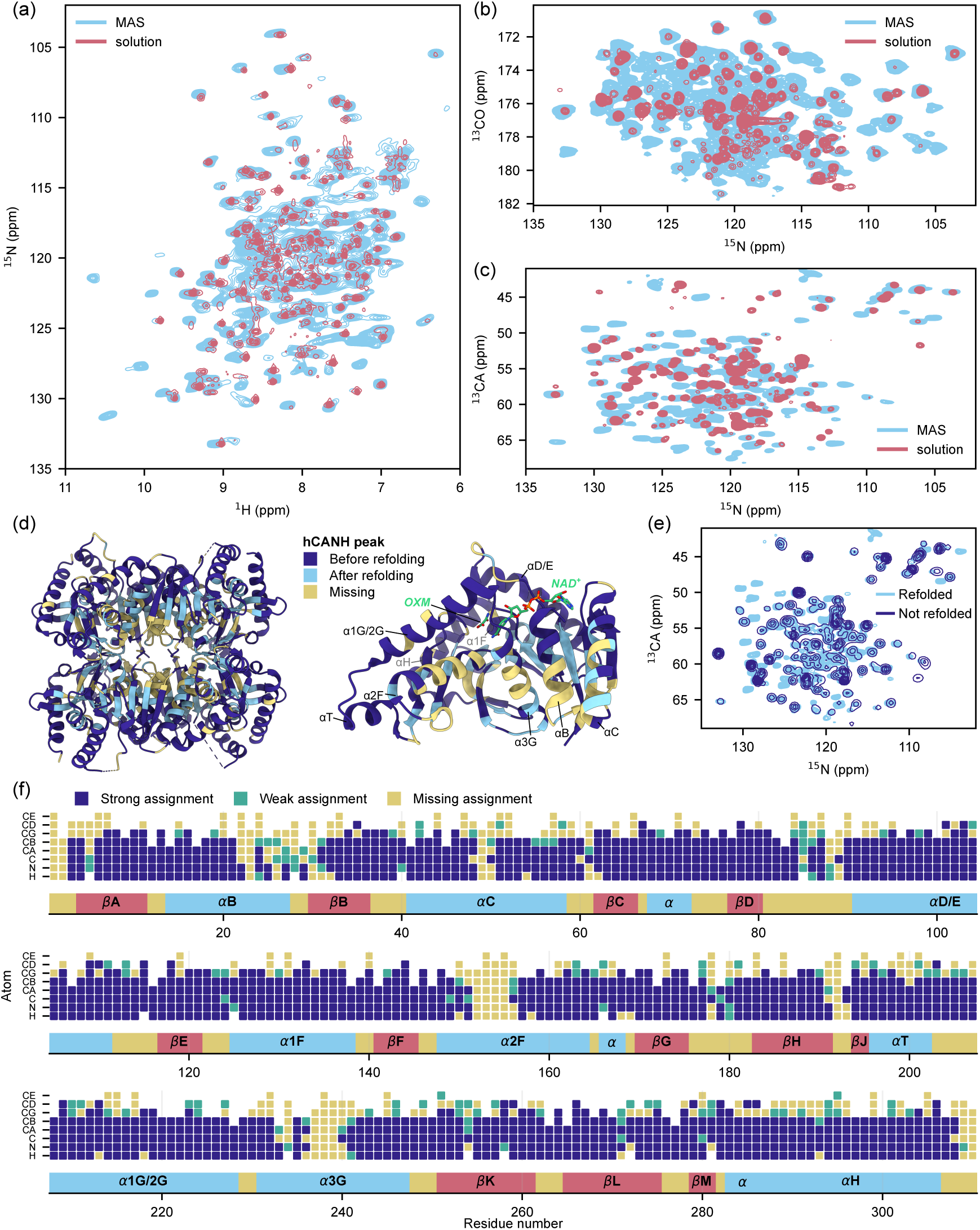
High-resolution solution– and MAS NMR of *Ii* MDH. (a) Overlay of solution– and MAS NMR 2D HN spectra. Despite the broader line-widths, the MAS NMR spectrum shows more peaks than the solution one. (b) Overlay of the N-CO projection of HNCO (solution) and hCONH (MAS) spectra. (c) Overlay of the N-CA projection of HNCA (solution) and hCANH (MAS). The difference in detectable peaks is exacerbated in 3D spectra, highlighting the limits of solution NMR for backbone assignment of large proteins. (d) Cartoon representation of the *Ii* MDH structure (PDB: 6QSS), highlighting residues that are visible before refolding, only after, or missing. Left: tetramer view; right: monomer view, with putative ligand binding sites – oxamate (OXM) and NAD^+^ aligned from *Thermus thermophilus* LDH, PDB: 2V7P. (e) Overlay of the hCANH spectra before and after refolding. Different contours are plotted for visual clarity. (f) Extent of MAS NMR assignment. Weak assignments are supported by a single peak, strong assignments by two or more. Extended version and statistics are provided in Suppl. Fig. S2.

### Assignment of backbone and side-chain heavy atoms from MAS NMR

Our assignment approach followed two complementary strategies, using either ^13^C-detected experiments with U-^13^C,^15^N labeled protein (4D CANCOCX and CONCACX, recorded with a CPMAS CryoProbe^37^; projections are provided in Figure S1) or amide-^1^H-detected experiments with the U-^2^H,^13^C,^15^N sample (3D hCANH, hCONH, hcaCBcaNH, hNcocaNH and 4D hcaCBCANH, hcaCBcaCONH, hCACONH, hCOCANH, hNCAcoNH). For the case of deuterated protein, we observed that the reprotonation of amide sites was incomplete, as commonly observed for highly stable proteins^38–41^. We used an unfolding/refolding procedure to exchange also those amide sites that are buried in the folded state, which made 59 additional signals observable (Figure 1d-e; for experimental details see Methods). Collectively, the complementary ^1^H– and ^13^C-detected experiments provided rich information for resonance assignment, with a total of 3849 manually picked cross-peaks for this 310-residue protein, listed in Table S1.

For the assignment we opted for an iterative procedure of the program FLYA^42^ – which uses a genetic algorithm for resonance assignment – and manual verification. This process turned out to be very efficient, and resulted in the assignment of 92 % of the backbone heavy atoms (C^a^, C’, N), 88 % of the amide hydrogens (H^N^), and 55 % of the side chain carbons (Figure 1f and extended in Figure S2). Overall, we obtained robust assignments – i.e., at least three out of four backbone atoms could be assigned – for 276 out of the 310 residues; the sequence stretches with missing or weak assignments include termini and loop regions (such as residues 1-2, 88-89, 191-193, 308-310), which is an often reported observation^28,36,43,44^, related to fast local dynamics that renders the dipolar-coupling based transfers less efficient^45^. More interestingly, a continuous structural segment that extends from the tetramer center to helix a2F (which bears the substrate-complexing residue Arg154) also resisted the assignment efforts (Figure 1d). This observation points to conformational dynamics on the µs-ms time scale; the investigation of this process is deferred to a dedicated manuscript. The residue-wise secondary structure positions, derived from these assigned chemical shifts using the TALOS-N approach^46^, are in excellent agreement with those found in the crystal structure (Figure S3).

With this near-complete backbone assignment from the sediment, we used two approaches for transferring the assignments to the solution NMR sample: matches of frequency triples in TROSY-based HNCO and HNCA backbone-assignment experiments – the only ones that provided a decent number of cross-peaks in solution – allowed identifying the majority of amide sites observed in the solution NMR TROSY spectrum; furthermore, a nuclear Overhauser effect spectrum (NOESY) connecting the amide sites confirmed and extended these assignments (Figure S4).

### Leveraging MAS-NMR assignments for complete assignment of methyl groups in solution

We aimed to obtain a comprehensive assignment of Met, Ile, Thr, Val and Leu methyl sites in solution, with the goal to subsequently probe dynamics and interactions. The near-complete backbone and substantial side-chain ^13^C assignment from MAS NMR described above allows for direct spectroscopic methods for methyl ^1^H-^13^C assignment. We combined several solution– and MAS NMR experiments, and studied their relative contributions to the assignment, as described in the following. In addition to these purely spectroscopic approaches, we prepared four single-point mutants (V168A, V27A, I50V, T152S), out of a total of 115 methyl-harboring residues.

We compared five spectroscopic approaches – two through-bond experiments in solution, one through-bond in the solid state, and two NOESY-based – differing in labeling scheme, sensitivity and residue coverage. The first route employed solution NMR, and a recently proposed isotope-labeling pattern of α-ketoisovalerate which results in the incorporation of ^13^C in the two methyl groups (γ1, γ2 in Val; δ1, δ2 in Leu), and at the α (Leu and Val) and β (Leu) positions; the carbon next to the methyls is ^12^C, and therefore the methyl carbons do not have one-bond ^13^C-^13^C couplings, which results in well-resolved singlet peaks. A specifically designed pulse sequence in solution connects both methyl groups to the C^α^(Val) and C^α^/C^β^(Leu) frequencies^47^. Because the latter frequencies are available in our case from the MAS NMR assignment, this approach immediately identifies both methyl groups for residues with known C^α^/C^β^assignments (Figure 2a, lines labeled HMBC-CC-HMQC; for this experiment, the V168A mutant was used). Using this approach allowed the assignment of 83 (71.6 %) Val and Leu methyls. The entire assigned spectrum is shown in Figure S5.

**Figure 2:**
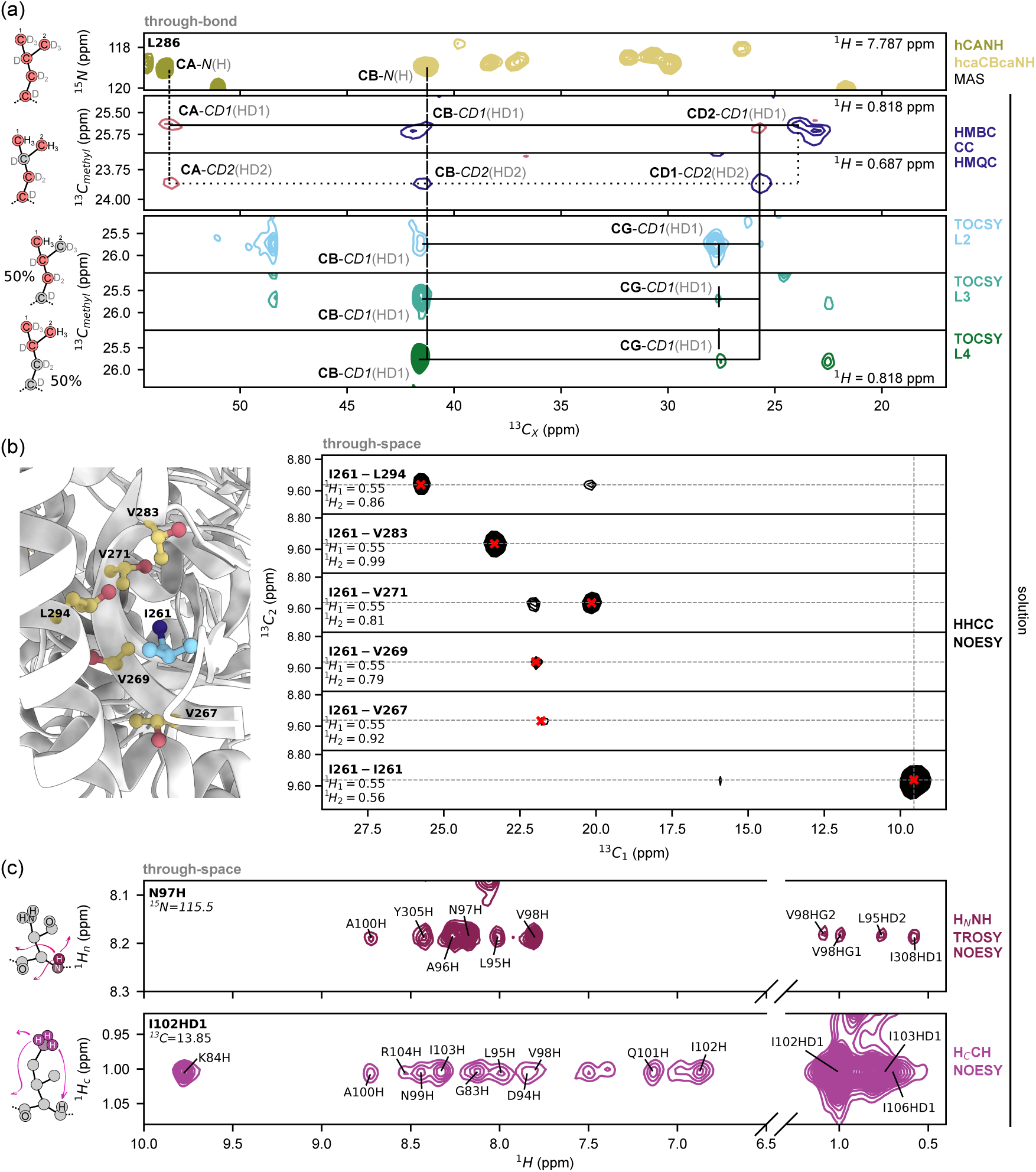
Solution NMR methyl assignment enabled by MAS-NMR derived assignment. (a) Through-bond experiments for Leu286 side-chain assignment in MAS and solution NMR. The different labeling schemes used, reported on the left, induce mild isotope shift and changes in peaks positions. From top to bottom: MAS NMR hCANH and hcaCBcaNH, solution NMR HMBC-CC-HMQC of a sample labeled with a-ketoisovalerate^47^ (both the slices corresponding to the two methyl groups are shown), solution NMR out-and-back TOCSY of a sample labeled with ketobutyrate and acetolactate precursors^48^ (three TOCSY mixing times – L2, L3, L4 – resulted in different extent of transfer, favoring directly bonded or further carbons). (b) Through-space methyl-methyl contacts of Ile261 detected in a solution NMR 4D H-H-C-C NOESY. The corresponding region in the crystal structure (PDB: 6QSS) is provided on the left. (c) Through-space backbone-methyl (top, H^N^-(TROSY)-N-H NOESY, Asn97) and methyl-backbone (bottom, H^C^-C-H NOESY, Ile102) contacts by solution NMR.

A second approach employed a labeling pattern that results in two kinds of molecules with linear ^13^C spin systems, and deuteration everywhere except at a single methyl group^48^: one pattern connects the methyl to the backbone (Val: γ1 *→* β*→* α; Leu: δ1 *→* γ*→* β; Ile: δ1 *→* γ1 *→* β*→* α), while the other connects the two methyls to each other (Val: γ2 *→* β*→* γ1; Leu δ2 *→* γ*→* δ1). Using a ^13^C-TOCSY based pulse sequence in solution allowed connecting the methyl to the ^13^C frequencies along the side chain up to C^a^ (Val, Ile) or C^A^ (Leu, Figure 2a, lines labeled TOCSY). The duration of the TOCSY mixing resulted in different extent of transfer, favoring either the directly bonded or further remote carbons (labeled L2, L3, L4 in Figure 2a and Figure S6; see Methods for details). Compared to the labeling pattern with alternate ^13^C spins described above, this experiment suffers from one-bond ^13^C-^13^C *J* couplings, resulting in broader ^13^C lines (note that virtual homo-decoupling is able to eliminate these splitting, but was not employed here). This second approach resulted in 72 Ile, Val and Leu methyls assigned (48.6 %). Overall, these TOCSY experiments performed worse than the HMCB-CC-HMQC at assigning Val and Leu residues – especially Val-γ2 and Leu-δ2 groups – but they nicely complemented it with the Ile assignments (which could not be obtained with the a-ketoisovalerate-labeled sample). The combination of the two strategies allowed the assignment of 117 Ile, Leu and Val methyl groups (79.1%, Figure S7).

Thirdly, we implemented MAS NMR experiments to connect the methyl to the rest of the side chain, in a 3D H^methyl^-C^methyl^-C correlation experiment, or to the backbone amide, in a C^methyl^-N^amide^-H^amide^ experiment, respectively (see Figure S8). The former experiment (hCCH) yielded 98 assigned cross-peaks (44 methyl groups, 29.7%), and is, in principle, a valuable strategy for methyl assignment. The latter experiment (hCccNH) had lower sensitivity and resulted in only 22 assignable peaks, presumably because the magnetization is spread along the side chain carbons, and the final ^13^C-^15^N step primarily selects the magnetization at the C^a^. Thus, while the connection of the methyl to the backbone is also possible in the solid state, we found here the solution-NMR approaches described above advantageous: not only the transfer efficiency is higher, but also the resolution of the ^1^H-^13^C spectrum is superior.

The experiments described above are not able to report on methyls of Thr and Met. As a fourth approach, we used several through-space NOESY experiments to cover also these, and to further support and extend the assignments obtained with the through-bond correlation experiments. A 4D methyl NOESY (Figure S10) of an ILV-labeled sample, exemplified in Figure 2b for I261, was found particularly powerful to generate unambiguous through-space connectivities, and complemented by a 3D H-H-C NOESY and a 3D H-C-C of a MIT-labeled sample (Figure S11). The combination of these two 3D experiments contributes to overcoming the assignment ambiguity, and it should be noted that their sensitivity is clearly superior to the one of the 4D, in particular for the H-H-C spectrum. The three spectra yielded a total of 295 non-diagonal peaks, resulting in the assignment of 99 Ile, Val, Leu, Met, Thr methyl groups (57.2 %)

A last approach leveraged the amide H-N assignments to connect methyl groups to the backbone via two complementary experiments: an amide-resolved H^N^-(TROSY)-N-H NOESY experiment^49^, and a methyl-resolved H^C^-C-H NOESY counterpart (Figure 2c and Figures S4,S9). Due to the high degeneracy of amide and methyl ^1^H frequencies, these spectra are laborious to work with, but they nevertheless allowed the assignment of 70 Ile, Leu and Val methyl groups (47.3%).

Collectively, these experiments – and the four single-point mutants that were required to resolve am-biguities – enabled the assignment of 100 % of Ile-δ_1_, Leu-δ_1_/-δ_2_, Val-γ_2_, Met– and Thr-γ and 63.4% of Val-γ_1_ (Figure S12). The extensive data set also allows us to evaluate the relative importance of the differ-ent approaches (summarized in Table S1). Solution NMR proved more efficient than MAS experiment for methyl assignment, largely due to the better resolved methyl spectra. The HMBC-CC-HMQC experiment was particularly powerful for Leu/Val assignments (both geminal methyls), while the TOCSYs allowed the assignment of majority of Ile. NOESY spectra completed the picture and were the only spectroscopic source of assignment for Met and Thr residues: while four-dimensional experiments remain preferable to obtain unambiguous assignment of contacts, three-dimensional HHC, HHN and HCC NOESY spectra allow greater sensitivity, and can represent a practical alternative when used in combination.

### Backbone dynamics of key functional regions identified by MAS NMR

Spin relaxation experiments in MAS NMR are sensitive to internal protein dynamics on a wide range of time scales^50,51^. While in solution, the overall tumbling dominates spin relaxation, and the site-to-site variations are generally very small, in the solid state spin-relaxation rate constants across the protein frequently differ by more than one order of magnitude. We focused on ^15^N rotating-frame relaxation (*R*_1_*_p_*), which is particularly sensitive to µs time-scale dynamics^52–54^. Given the large size of the protein, we employed a pseudo-4D experiment consisting of a series of 3D hCANH spectra to unambiguously quantify the dynamics of as many residues as possible; the high sensitivity of this experiment with a 1.9 mm rotor, combined with non-uniform sampling, allowed recording a full relaxation series of 7 spectra within 4 days. Figure 3a,c shows the *R*_1_*_p_* rate constants in sedimented *Ii* MDH, which highlights well-defined regions with pronounced µs mobility. Relaxation decays are presented in Figure S13. The so-called catalytic loop, which harbors the catalytic histidine His178, is one of these regions; interestingly, a three-residue stretch comprising His178 itself was not detectable in these relaxation experiments; their low intensity is potentially due to the strong relaxation induced by µs dynamics. Residues 85-92, forming the “mobile surface loop”, also feature enhanced *R*_1_*_p_* relaxation; this part is supposed to close upon ligand binding, as shown by previous crystal structures of homologs^55,56^. Our data show that this region undergoes extended µs dynamics already in the apo state. The region with the most pronounced µs dynamics is the so-called Mobile Region 2 (MR2), which connects helices aT and a1G/2G. This region is associated with structural reorganizations between active and inactive states in allosteric lactate dehydrogenases^57,58^, and is not expected to do the same in non-allosteric malate dehydrogenases. Together with the observation of unassigned regions in a3G and a2F, harboring the substrate-chelating Arg154, this data suggests that the active site of *Ii* MDH explores multiple conformations and is not in a static activated one. Lastly, elevated *R*_1_*_p_* rates are found for the N– and C-termini, which are expected to be flexible.

**Figure 3:**
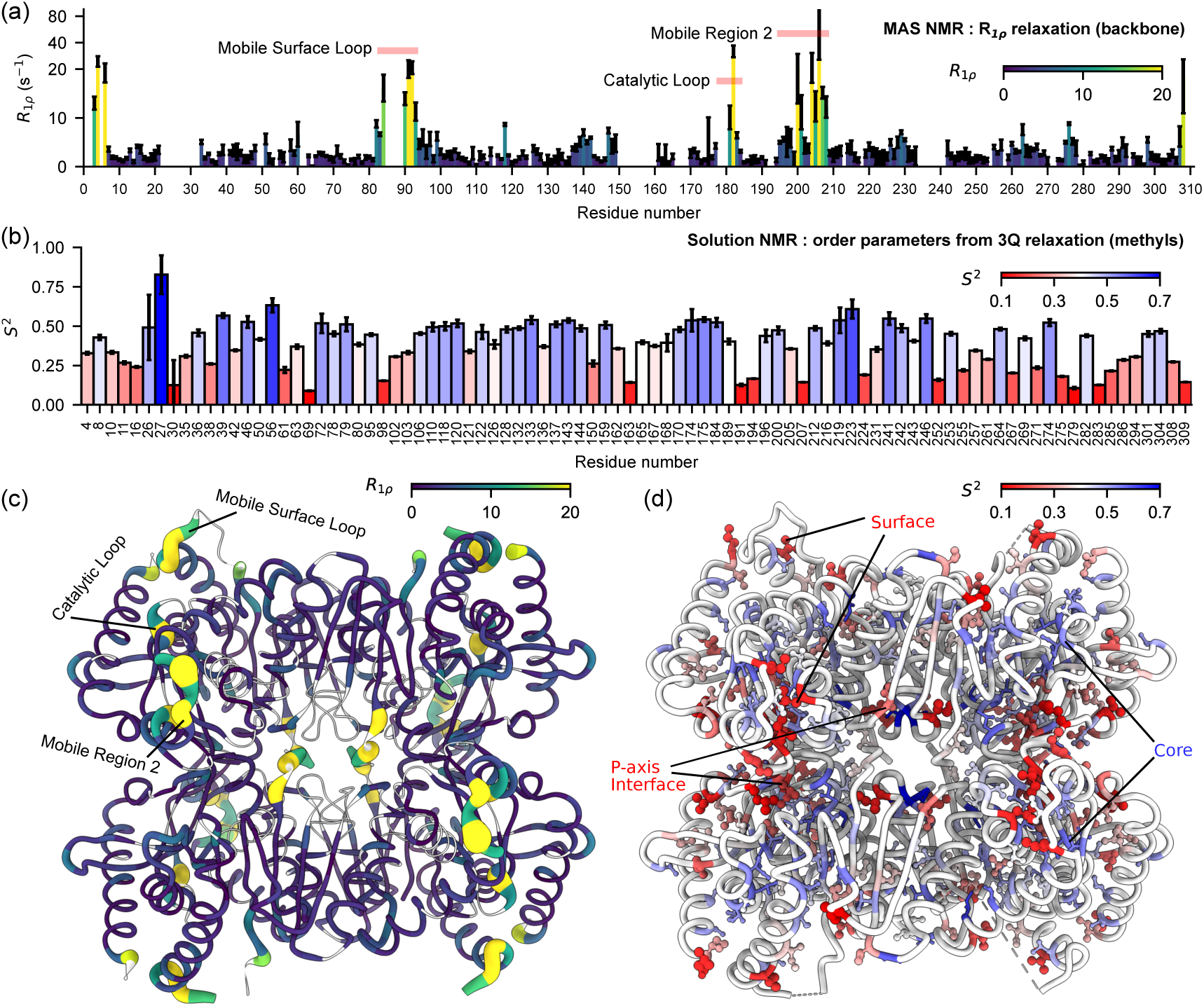
Backbone-directed MAS NMR and methyl solution NMR site-specific insights into the dynamics of *Ii* MDH. (a, c) ^15^N *R*_1_*_p_* rate constants acquired at 38 kHz MAS frequency and a spin-lock radio-frequency (RF) field strength of 10 kHz. (b, d) Methyl-axis order parameters, *S*^2^, determined with a triple-quantum relaxation violated coherence transfer experiment^59^.

Collectively, backbone dynamics measurements clearly identify µs mobility in regions of key functional importance.

### Methyl-detected dynamics in solution highlight dynamic residues, including in the core

We studied side chain dynamics to gain complementary views of *Ii* MDH’s mobility. Not only do methyls sense different types of motions, but in the case of *Ii* MDH, methyls also expand the accessible structural parts: in some parts where the backbone remained unassigned, methyl probes are available (residues 26, 27, 30, 150, 159, 241, 242). Solution NMR methyl spectra are advantageous in terms of resolution for *Ii* MDH, which prompted us to use solution NMR triple-quantum relaxation experiments^59^ to quantify the order parameters of methyls, 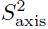. This parameter describes the motional restriction of the methyl axis, ranging from 1 (restricted) to 0 (unrestricted motion), on time scales shorter than the overall tumbling (ca. 40 ns).

The obtained order parameters of Ile, Leu and Val in *Ii* MDH follow expected overall trends, where surface-exposed methyl groups are more disordered than those in the hydrophobic core (Figures 3b,d, and S14). However, there are notable exceptions that are deeply buried yet show high flexibility, such as Ile30, Ile61, Leu163, Ile191, Ile252, Ile275 and Leu279. Interestingly, these involve the point of contact between MR2 and Mobile Region 1 (MR1, at the end of helix a2F^57^) and several residues along the P-axis dimer-dimer interface, suggesting rather loose packing of these hydrophobic clusters.

Low methyl-axis order parameters reflect the mobility of the backbone as well as the transitions between rotamer states, i.e. rotations around the side chain dihedral angles *χ*_1_ and *χ*_2_. Chemical shifts provide an orthogonal view on the population of rotamer states^60–63^. Given the extensive assignment we have obtained, including both Val/Leu *pro*S and *pro*R methyls, quantitative estimates of rotamer-state populations are readily available from our data (Figure S15). Interestingly, we observed a mild correlation between trans/gauche-/gauche+ populations and 3Q relaxation-derived *S*^2^ for Leu (R=0.677) and Ile (R=0.580) residues, suggesting that rotamer equilibrium can partially contribute to the order parameters. On the other hand, we found only poor correlation for Val (R=0.262) residues. Three factors could contribute to this: first, the timescales probed by chemical shifts (up to ms) and relaxation (≲40 ns) substantially differ. Consequently, excess motion seen by chemical shifts may occur on the time scale from tens of ns to ms. Second, estimating Val rotamer populations is complicated, and it relies on maximizing the agreement to quantum chemical calculations using a genetic algorithm^60^. Third, for Val in particular, its shorter side-chain could cause the methyl groups to experience more of the backbone dynamics, and therefore rotamer equilibrium to contribute less to the measured order parameters.

### Backbone-observed solution NMR for quantitative ligand-binding studies

Solution methods are clearly at their best for probing inter-molecular interactions. Following chemical-shift perturbations (CSPs) upon addition of substrate is a powerful way to not only identify the residues involved in the binding, but also the interaction strength. However, methyl-directed experiments fall short of seeing interactions at polar residues, and therefore often miss the residues directly involved in the active sites. We used H-N TROSY-HSQC experiments to probe the binding of oxamate, a structural analogue of pyruvate and known competitive inhibitor^64,65^. Using two-dimensional lineshape analysis in NMR-TITAN^66^ we determined a *K_d_* of 22.5 *±* 1.4 mM (Figure S16).

Figure 4 shows spectra and residue-specific CSPs upon addition of oxamate to a solution of *Ii* MDH. It should be mentioned that the solution NMR backbone spectrum exhibits fewer peaks than the MAS NMR one, especially missing residues in the surrounding of the non-assigned regions, possibly due to conformational-exchange broadening. Some of the residues that are most affected by binding are in the catalytic loop harbouring the catalytic His178, i.e. the loop which we showed to undergo µs motions (to be noted, no methyl-harboring residue is present in this loop). Similarly, Thr230 – another residue that makes direct contact to the substrate, on Mobile Region 3 (MR3) between helices a1G/2G and a3G^67^ – also shows large CSPs.

**Figure 4:**
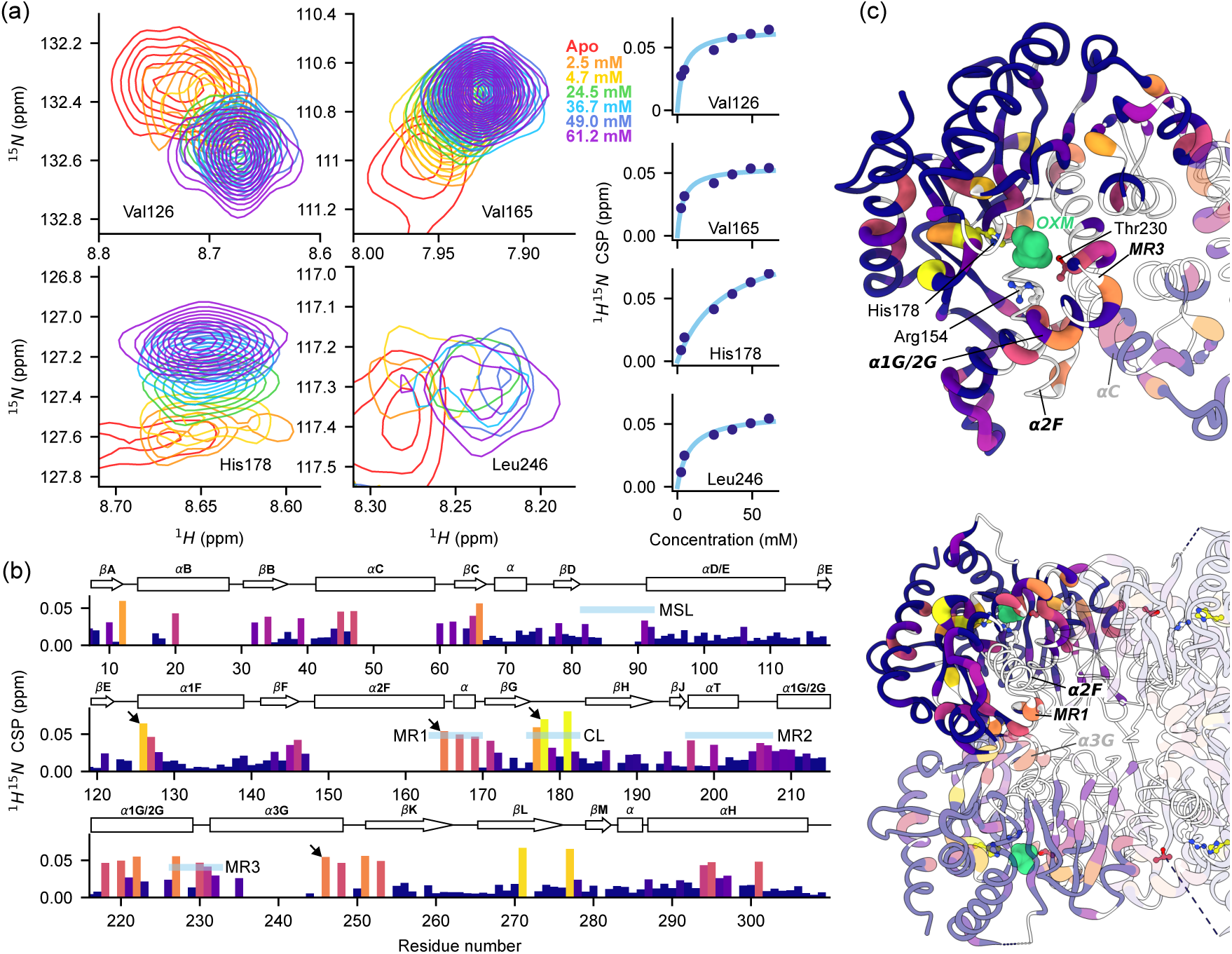
Backbone-observed oxamate binding in *Ii* MDH. (a) Concentration-dependent CSPs upon oxamate binding for selected residues in *Ii* MDH. Left: overlay of 2D NH spectra recorded throughout the titration. Right: CSPs as a function of oxamate concentration. (b) Barplot of CSPs in the presence of 61.2 mM oxamate. Large perturbations are observed for the catalytic loop (CL), MR1, MR3 and helix a3G. No significant changes are found at the mobile surface loop (MSL) and helix aD/E. Residues shown in panel a are highlighted with arrows. (c) Worm and color gradient representation of CSPs onto the *Ii* MDH structure (PDB: 6QSS). Top: top view; bottom: front view. The subunits are shown in different transparency for visual clarity.

While these effects are expected — given that these residues directly form the substrate-binding site — our data points to an interesting and less expected region: strong CSPs for Thr146, Thr148, Val165, Ile167, Ala169 and Asp171 show that helix a2F and MR1 undergo a significant structural rearrangement. This helix bears the residue Arg154, which is known to bind substrate. Importantly, the Arg154-substrate interaction and the rearrangement of helix a2F/MR1 have been shown to be crucially involved in allosteric signal transmission^6,57,68^. Moreover, the CSP data point to long-range structural modifications also in a3G: while on the one side of this helix (Gln227, Thr230) the observed CSPs can be ascribed to direct interaction with oxamate, large CSPs are also seen for Gln246, Asn248, located at its other extremity, suggesting a rearrangement of the entire helix. Interestingly, the end of helix a3G of one subunit is in direct contact with the MR1 of the P-axis related subunit, creating a connection between active sites reminiscent of allosteric LDHs^69^. As noted above, *Ii* MDH lies phylogenetically in between the non-allosteric MDH3 and allosteric LDH groups, and has been proposed as a pre-allosteric enzyme^8^. Notably, this interface is surrounded by residues with low order parameter (see previous section) which could allow motion between different conformations without engaging in strong hydrophobic interactions.

Interestingly, we did not observe any significant perturbation for helix aD/E and the mobile surface loop, indicating that this loop remains open in the presence of oxamate – in contrast to the interaction with the actual substrate, oxaloacetate, which leads to loop closure. We ascribe this observation to the fact that the substrate analogue oxamate lacks the C2 ketone of oxaloacetate that engages Arg92 and triggers loop closure^55^ (Arg86 will subsequently interact with the C4 carboxylate). Of note, the fact that our data highlight the absence of loop closure with oxamate suggests that structures solved in the presence of oxamate and not the native substrates might not reflect the enzyme-substrate complex correctly. A thorough investigation of binding with different substrates is deferred to a dedicated manuscript.

Collectively, the availability of assigned backbone N-H resonances with N-H TROSY — which is still feasible for proteins up to hundreds of kDa — allowed probing substrate interactions, including polar residues that would escape detection by methyl-TROSY NMR.

## Conclusions

Determining the functional mechanisms of enzymes requires going beyond static three-dimensional structures to characterize the dynamics and ligand interactions that underlie catalysis and allostery. For large oligomeric proteins, this information has been difficult to obtain by NMR: de novo backbone assignment fails in solution at high molecular weight, methyl-TROSY approaches leave polar residues invisible, and the conventional mutagenesis-based route to methyl assignment is laborious. The present work shows that integrating MAS NMR of sedimented samples with solution NMR resolves these bottlenecks in a practical and largely automated manner, and opens a comprehensive view of structure, dynamics and interactions in a 134 kDa tetrameric enzyme.

The foundation of our approach is the backbone resonance assignment, which is achievable in the sedimented sample by MAS NMR but not in solution at this molecular weight. Near-complete assignment of backbone heavy atoms and amide protons was obtained by combining ^1^H– and ^13^C-detected high-dimensional experiments with iterative automated assignment using FLYA. Transferring these assignments to the solution-state sample through direct spectroscopic connections to the known backbone frequencies, yielded 91.3% coverage of Ile, Leu, Val, Met and Thr methyls — a level of completeness that would be difficult to achieve by mutagenesis alone for a protein of this size. Residues that resisted assignment are predominantly located in loop regions or at the tetrameric interface including secondary structures. The fact that these gaps cluster in regions of pronounced µs dynamics suggests that conformational exchange, rather than spectral overlap, is responsible: missing resonances thus carry functional information in their own right.

With this assignment in hand, MAS and solution NMR each contributed distinct and complementary dynamical information. ^15^N *R*_1_*_p_* relaxation measurements in the sedimented sample – resolving 244 individual sites in a series of 3D relaxation measurements – revealed pronounced microsecond-timescale backbone mobility in three functionally important regions: the catalytic loop harbouring His178, the mobile surface loop (residues 85–92) that is expected to close upon substrate binding, and Mobile Region 2. Methyl-axis order parameters from solution NMR provided a complementary picture of side-chain dynamics, identifying additional flexible sites in the hydrophobic core — including deeply buried residues such as Ile30, Ile61, Leu163, Ile191, Ile252, Ile275 and Leu279 — that are not captured by backbone measurements alone. The two sets of dynamics measurements are thus not redundant: backbone *R*_1_*_p_* is uniquely sensitive to microsecond motions and reports on the peptide plane, while methyl order parameters integrate pico-to-nanosecond backbone and side-chain dihedral fluctuations.

The substrate analogue oxamate produced chemical shift perturbations across a wide network of backbone amide sites, detectable by ^1^H-^15^N TROSY in solution. Beyond the expected contacts at the active site — the catalytic loop and Thr230 — significant perturbations were observed at helix *ε*2F, which bears the substrate-chelating Arg154, and propagate to the distal end of helix *ε*3G, suggesting a concerted rearrangement of structural elements known to be involved in allosteric signal transmission. These long-range effects would have been invisible using methyl-TROSY alone, underscoring the value of backbone-level observation for mapping interaction networks in large proteins.

Taken together, these results establish a practical workflow for the NMR study of large oligomeric assemblies that integrates the strengths of both approaches: sedimented-sample MAS NMR for backbone assignment and microsecond dynamics, and solution NMR for side-chain dynamics and ligand interactions. The workflow is efficient — relying on automated assignment tools and a limited number of isotope-labeling schemes — and yields an unusually complete set of probes distributed across backbone and side chains of five amino acid types. We anticipate that this strategy will be broadly applicable to other large oligomeric proteins for which methyl-TROSY alone provides insufficient coverage, and that it will be particularly powerful in cases, such as the MDH/LDH superfamily, where understanding allostery requires simultaneous access to dynamics at the active site, the subunit interface, and the regions connecting them.

## Materials and Methods

### Protein production

A DNA insert coding for the sequence of *Ignicoccus islandicus* malate dehydrogenase (uniprot ID A0A0U3FQH7) was cloned into a pET-20b vector between the NdeI and BamHI cloning sites (codon-optimized, obtained from GeneCust, https://genecust.org). Protein production was performed in *E. coli* BL21 Star (DE3) in suitably prepared media for isotope labeling, as follows. For the production of fully protonated protein (used for carbon-detected experiments) the cells were grown in 100% H_2_O M9 medium, supplemented with 1 g L^-1^ ^15^N ammonium chloride and, 2 g L^-1^ ^13^C_6_ glucose. For the production of deuterated proteins, cells were adapted to D_2_O in gradual steps (0, 50, 100 %) of 8h or over night duration. All cultures were supplemented with 1 g L^-1^ ^15^N ammonium chloride and 2 g L^-1^ glucose which was either ^13^C,1,2,3,4,5,6,6-^2^H_7_ labeled (MAS NMR samples and a-ketoisovalerate labeling) or 1,2,3,4,5,6,6-^2^H_7_ labeled (all the other side-chain labeling schemes).

For producing the sample that was used for the HMBC-CC-HMQC approach, we used (2-^13^C, 3-methyl-^13^C, 4-^13^C, 3-^2^H) a-ketoisovalerate^47^ (Mag-Lab, https://www.mag-lab.eu, compound L15), which was added to the culture 1 hour prior to induction (150 mg L^-1^).

For the samples with linearized ^13^C chains, used for the TOCSY experiments in solution and in solids, we followed the protocol by Kerfah *et al.*^48^ with some modifications: no alanine precursor was used; the glucose was deuterated, but not ^13^C labeled, which means that the a position of Leu is not ^13^C labeled in the final protein. We purchased linearized precursors for side-chain labeling in form of a precursor kit (NMR-bio, https://nmr-bio.com). Acetolactate and ketobutyrate precursors were added 1 hour or 20 minutes before induction, respectively.

For the single-methyl labeling of Ile, Val, Leu (ILV sample) or Ile, Met and Thr (MIT sample), used for NOESY experiments, we purchased TLAM-Iδ_1_LV-proS and TLAM-Iδ_1_M Tγ precursor kits (NMR-bio) and followed the manufacturer’s instructions.

Cultures were started at an OD_600_ of 0.2 and grown at 37 *^→^*C. Expression was induced when OD_600_ reached 0.6-0.8 and performed at 37 *^→^*C for 3-5 hours or at 20 *^→^*C over night.

Cells were harvested by centrifugation (15 min, 4000 rcf), pellets were resuspended in Buffer L and lysed either by sonication, using a Q700 ultrasonic processor (Qsonica, 40% amplitued, 6 mins operating time) or high pressure (continuous high pressure cell disruptor, 25-30 kpsi, 2 cycles). Initial purification was performed by heat shock of the lysates (75*^→^* C, 20 min) before high-speed centrifugation (50000 rcf, 60 min, 4*^→^* C). The supernatant was filtered (0.22 µm pore size) before applying it on an anion exchange Resource™ Q column (GE Healthcare) pre-equilibrated with Buffer A. Protein was eluted with a linear gradient of Buffer B (10 column volumes, 60 ml). Protein containing fractions were concentrated to <2 ml and applied on a Size Exclusion Chromatography HiLoad 26/600 Superdex 200 PG column (Sigma-Aldrich), pre-equilibrated with Buffer A.

For the unfolding/refolding procedure, the protein was diluted to 0.1 mg/ml in 6 M Guanidinium chloride and incubated at 4 *^→^*C over night. Afterwards, unfolding was achieved by heat shock (85*^→^*C, 1 hour). The sample was allowed to cool down before dialysis to Buffer A in two steps. Lastly, the sample was centrifuged (20000 rcf, 20 min) to remove aggregates, and concentrated back in Buffer A.

**Table 1:** Buffers compositions.

|  |  |
| --- | --- |
| Buffer L | 50 mM NaCl, 50 mM Tris-HCl, 0.025 mg/ml RNase, 0.05 mg/ml DNase,<br>2 mM MgCl<sub>2</sub>, protease inhibitor (cOmplete™, EDTA-free), pH 7 |
| Buffer A | 50 mM NaCl, 50 mM Tris-HCl, pH 7 |
| Buffer B | 1 M NaCl, 50 mM Tris-HCl, pH 7 |

### MAS NMR spectroscopy

Samples for MAS NMR were prepared by ultra-centrifugation of the protein solution (at ca. 15 mg/mL, H_2_O-based Buffer A) into MAS NMR rotors using an in-house built ultracentrifugation tool adapted for a Beckman SW32 rotor: a 1.9 mm Bruker BioSpin rotor (part number H123832) was used for ^1^H-detected experiments (assignment, dynamics), and a 3.2 mm rotor for the CPMAS CryoProbe (volume 81 µL with an insert for wet samples) for the ^13^C-detected assignment experiments. The rotors were filled by over-night centrifugation at 20,000 rpm (68300 g). Rotor caps were glued with two-component epoxy glue to avoid dehydration.

All ^1^H-detected MAS NMR experiments were recorded on a Bruker BioSpin NEO spectrometer operating at a ^1^H frequency of 700 MHz equipped with a 1.9 mm HXY MAS probe tuned to ^1^H, ^13^C and ^15^N frequencies. All ^13^C-detected experiments were recorded on a 600 MHz Bruker NEO spectrometer equipped with a CPMAS CryoProbe^37^. Experimental times, MAS speeds and sample labeling for the assignment spectra are reported in Table S3.

*R*_1_*_p_* relaxation was measured in a series of 7 three-dimensional hCANH spectra with non-uniform sampling for a total duration of 4 days. The spinlock frequency was set to 10 kHz and the relaxation delays were 5, 20, 40, 60, 80, 100 and 130 ms. Relaxation rates were determined by fitting a mono-exponential function to the peak intensity decays.

### Solution-state NMR spectroscopy

All solution-state NMR samples were prepared in H_2_O– or D_2_O-based Buffer A, added with 3.75% D_2_O, 0.4 mM DSS to enable lock and spectra referencing. For the oxamate titration, pH was set to 6, in line with pyruvate activity tests^8^. Concentration of samples for solution-state NMR are indicated in Table S2. All spectra were obtained on a 800 MHz Avance NEO spectrometer equipped with a TCI ^1^H/^13^C/^15^N 5 mm cryoprobe.

Methyl-axis order parameters *S*^2^ were determined through a 3Q-relaxation experiment^59^, in two series of 8 two-dimensional CH experiments for a total duration of 109 hours. Relaxation delays of 2, 4, 6, 8, 11, 15, 20 and 25 ms were used. The overall-tumbling correlation time was determined through the TRACT cross-correlated relaxation experiment^70^. More details about these two experiments, fitting routines and equations, are provided in the Supplementary Information, and scripts for these fits are provided alongside this article on the publisher’s website.

## Associated content

Resonance assignments were deposited at the Biological Magnetic Resonance Data Bank (BMRB ID 53910).

## Supporting information

The following files are available free of charge.

- Supplementary_Information.pdf: description and equations for fitting TRACT and 3Q-relaxation experiments; Supplementary Figures S1-S16 including assignment spectra and statistics, pulse sequences, secondary structure and rotamer probabilities, fits of dynamics and ligand binding experiments; Tables S1-S3 including peaklists statistics and acquisition parameters of solution and MAS NMR experiments.
- Scripts_submission.zip: fit routines in python language of TRACT and 3Q-relaxation experiments.
- pulsesequences.zip: four pulse sequences and acquisition parameters: (i) hCCH and (ii)hCccNH TOCSY experiments in solids, and (iii) TOCSY in solution and (iv) HMBC-CC-HMQC.

## Acknowledgements

We thank Dominique Madern (IBS Grenoble) for many discussions and for providing the plasmid. We are grateful to Rime Kerfah and Elodie Crublet (NMR-bio), and Roman Lichtenecker (Univ. Vienna/Mag-Lab) for advice on the use of isotope-labeled compounds. This work has been funded in part by the Austrian Science Fund (project “AlloSpace. The emergence and mechanisms of allostery”, grant number 10.55776/I5812). This research was supported by the Scientific Service Units (SSU) of Institute of Science and Technology Austria (ISTA) through resources provided by the Nuclear Magnetic Resonance and the Lab Support Facilities. We thank Petra Rovó, Megha Mohan and Margarita Valhondo Falcón for excellent support of the NMR facility.

---

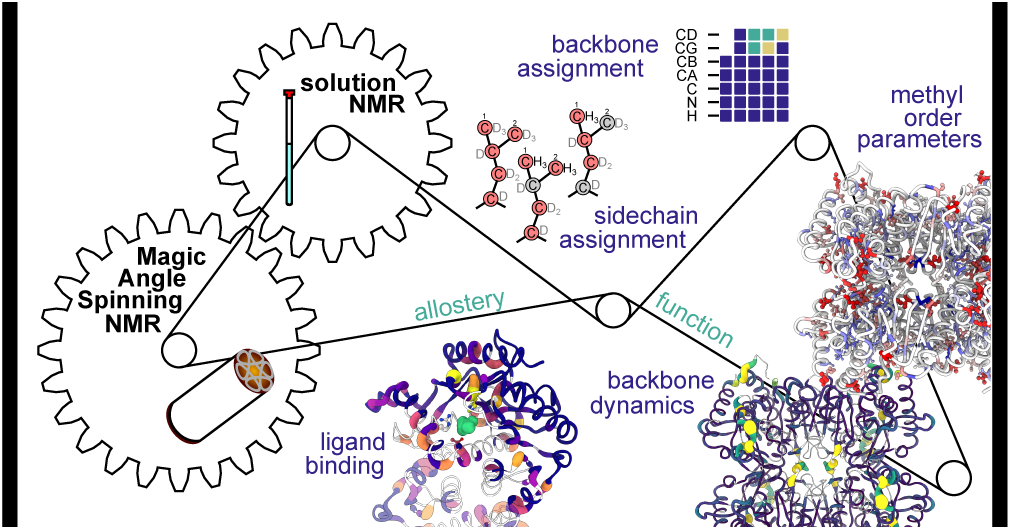

