## Supplementary Information for "Integrated solid/solution NMR assignment allows mapping dynamics and ligand binding in a 134 kDa enzyme"

### Determination of methyl order parameters and overall-tumbling correlation time

The methyl-axis order parameters were determined from a triple-quantum (3Q) relaxation violated coherence transfer NMR experiment in solution, described elsewhere[59]. In brief, the experiment measures the time-dependence of the build-up of forbidden 3Q coherence (with intensity $I_A$) and the intensity evolution of a single-quantum coherence with intensity $I_B$ as a function of the relaxation delay $t$. The ratio of these intensities depends, *inter alia* the sought cross-correlated relaxation with a rate $\eta$, and a term $\delta < 0$ that accounts for coupling between the rapidly and slowly decaying $^1$H SQ coherences due to relaxation with external protons, as follows:

$$I_A/I_B = \frac{3}{4}\frac{\eta \tanh\left(\sqrt{\eta^2+\delta^2}\,t\right)}{\sqrt{\eta^2+\delta^2} - \delta \tanh\left(\sqrt{\eta^2+\delta^2}\,t\right)} \tag{1}$$

$$\eta = \frac{9}{10}\left(\frac{\mu_0}{4\pi}\right)^2 [P_2(\cos\theta_{axis,HH})]^2 \frac{\gamma_H^4 \hbar^2}{r_{HH}^6} S^2 \tau_c \tag{2}$$

.

Two data sets, corresponding to $I_A$ and $I_B$ were collected and the experimental ratio was fitted using a standard chi-squares fit procedure, with $\eta$ and $\delta$ as fit parameters.

$$\chi^2 = \sum_i \left(\frac{I_{A/B}^{\text{model}}(t_i) - I_{A/B}^{\text{data}}(t_i)}{\sigma_i}\right)^2 \tag{3}$$

where $I_{A/B}$ is a shorthand notation for the intensity ratio. The constants are as follows:

$$\begin{array}{lll} \gamma_H = 2.67522 \times 10^8\ \mathrm{rad\,s^{-1}\,T^{-1}} & \gamma_C = 6.7283 \times 10^7\ \mathrm{rad\,s^{-1}\,T^{-1}} & \gamma_N = -2.7126 \times 10^7\ \mathrm{rad\,s^{-1}\,T^{-1}} \\ \mu_0 = 4\pi \times 10^{-7}\ \mathrm{T^2\,m^3\,J^{-1}} & h = 6.626 \times 10^{-34}\ \mathrm{J\,s} & \hbar = 1.054 \times 10^{-34}\ \mathrm{J\,s} \\ r_{HH} = 1.813 \times 10^{-10}\ \mathrm{m} & \theta_{axis,HH} = \frac{\pi}{2} & P_2 = \frac{1}{2}(3\cos^2\theta - 1) \end{array} \tag{4}$$

The 3Q relaxation experiment reports on the product of the methyl-axis order parameter and the overall-tumbling correlation time, $S^2_{axis}\tau_c$[59]. Assuming isotropic overall tumbling, the conversion from the fitted $S^2_{axis}\tau_c$ to $S^2_{axis}$ is trivial. To assess how well this isotropic-tumbling assumption is justified, we used the program HydroNMR[71] that models the diffusion properties from the structure, using the crystal structure (6QSS) as input.

The principal axes of the diffusion tensor, reported by HydroNMR are $D_z = 3.786 \times 10^6$ s$^{-1}$ (largest, i.e., fastest rotation), $D_x = 3.428 \times 10^6$ s$_{-1}$ (middle), $D_y = 3.098 \times 10_6$ s$^{-1}$ (smallest, slowest rotation). The anisotropy ratio is ca. 1.2 (the fastest axis tumbles ca. 20% faster than the slowest). This modest anisotropy shows that the assumption of a uniform value of $\tau_c$ does not lead to strong distortion of the order parameter profile.

In order to obtain an experimental estimate of the overall-tumbling correlation time we used the cross-correlated relaxation experiment known as TRACT[70], as implemented in NMRlib[72]. The decays of the two components of $^{15}$N coherence, $R_{2,\alpha}$ and $R_{2,\beta}$ were fitted with mono-exponential functions, using the bulk of the amide signals (see Figure S14).

$$C_{\text{CCR}} = S^2 \frac{4}{15} \frac{\mu_0}{4\pi} \hbar \gamma_N^2 \gamma_H B_0 \Delta\sigma \, r_{NH}^{-3} \tag{5}$$

$$\tau_c = \frac{|R_{2,\beta} - R_{2,\alpha}|}{2C_{\text{CCR}}} \tag{6}$$

The constants were:

$$r_{NH} = 1.021 \times 10^{-10} \text{ m} \qquad B_0 = 18.8 \text{ T} \qquad \Delta\sigma = 140 \times 10^{-6} \tag{7}$$

The fit routines in python language of both experiments are provided with the data repository; see main text for the access details.

### Supplementary tables

Table S1: Assignment spectra statistics.

| Backbone assignment experiment (MAS) | Number of peaks |
|---|---|
| hCANH (3D) | 250 |
| hCONH (3D) | 222 |
| hcaCBcaNH (3D) | 193 |
| hcaCBCANH (4D) | 223 |
| hCACONH (4D) | 177 |
| hCOCANH (4D) | 243 |
| hcaCBcaCONH (4D) | 430 |
| hNcocaNH (3D) | 220 |
| hNCAcoNH (4D) | 228 |
| CONCACX (4D, C-det.) | 864 |
| CANCOCX(4D, C-det) | 799 |

| Sidechain assignment experiment | Number of peaks | Assigned methyls |
|---|---|---|
| ILV HMBC-CC-HMQC | 180* | 83 (71.6%) |
| ILV TOCSYs (L2/L3/L4) | 90*/93*/116* | 72 (48.6%) |
| MAS ILV TOCSY | 102* | 44 (29.7%) |
| MAS ILV hCccNH TOCSY | 23 | |
| ILV HHCC NOESY | 105* | 70 (47.3%) |
| MIT HHC/HCC NOESY | 86*/104* | 55 (96.5%) |
| ILV $H^N$-(TROSY)-N-H NOESY/$H^C$CH NOESY | 315*/183* | 70 (47.3 %) |

* : Diagonal peaks were omitted from the count

Table S2: Acquisition parameters of solution NMR spectra.

| Sample labeling | Experiment | Concentration | Solvent | Exp. time |
|---|---|---|---|---|
| U-$^2$H,$^{13}$C,$^{15}$N, α-ketoisovalerate | HMBC-CC-HMQC | 400 µM | $D_2O$ | 63h |
| U-$^2$H,$^{15}$N, ketobutyrate, acetolactate | TOCSYs (L2, L3, L4) | 400 µM | $D_2O$ | 93h, 68h, 108h |
| U-$^2$H,$^{15}$N, ketobutyrate, acetolactate | $H^N$-N-H NOESY/$H^C$CH NOESY | 400 µM | $H_2O$ | 69h, 73h |
| U-$^2$H,$^{15}$N, Ile-$\delta_1$, Leu-$\delta_2$, Val-$\gamma_2$ | HHCC NOESY | 400 µM | $D_2O$ | 212h |
| U-$^2$H,$^{15}$N, Ile-$\delta_1$, Leu-$\delta_2$, Val-$\gamma_2$ | TRACT, 3Q-relaxation | 400 µM | $H_2O$ | 3h, 110h |
| U-$^2$H,$^{15}$N, Ile-$\delta_1$, Met-ε, Thr-γ | HHC NOESY, HCC NOESY | 400 µM | $D_2O$ | 55h, 77h |
| U-$^2$H,$^{13}$C,$^{15}$N (refolded) | $^1$H-$^{15}$N-TROSY | 496.3 µM | $H_2O$ | 3h |
| U-$^2$H,$^{13}$C,$^{15}$N (refolded) | $^1$H-$^{15}$N-TROSY | 259.1 µM<br>OXM: 2.5 mM | $H_2O$ | 3h |
| U-$^2$H,$^{13}$C,$^{15}$N (refolded) | $^1$H-$^{15}$N-TROSY | 389.5 µM<br>OXM: 4.7 mM | $H_2O$ | 3h |
| U-$^2$H,$^{13}$C,$^{15}$N (refolded) | $^1$H-$^{15}$N-TROSY | 586.3 µM<br>OXM: 24.4 mM | $H_2O$ | 3h |
| U-$^2$H,$^{13}$C,$^{15}$N (refolded) | $^1$H-$^{15}$N-TROSY | 447.6 µM<br>OXM: 36.7 mM | $H_2O$ | 3h |
| U-$^2$H,$^{13}$C,$^{15}$N (refolded) | $^1$H-$^{15}$N-TROSY | 442.0 µM<br>OXM: 49.0 mM | $H_2O$ | 3h |
| U-$^2$H,$^{13}$C,$^{15}$N (refolded) | $^1$H-$^{15}$N-TROSY | 395.7 µM<br>OXM: 61.2 mM | $H_2O$ | 3h |

Table S3: Acquisition parameters of MAS NMR spectra.

| labeling | Experiment | Rotor size (mm) | MAS (kHz) | Exp. time |
|---|---|---|---|---|
| U-$^2$H,$^{13}$C,$^{15}$N (refolded) | hCANH | 1.9 | 38 | 94h |
| U-$^2$H,$^{13}$C,$^{15}$N (refolded) | hCONH | 1.9 | 38 | 50h |
| U-$^2$H,$^{13}$C,$^{15}$N | hcaCBcaNH | 1.9 | 38 | 72h |
| U-$^2$H,$^{13}$C,$^{15}$N (refolded) | hcaCBCANH | 1.9 | 38 | 165h |
| U-$^2$H,$^{13}$C,$^{15}$N (refolded) | hCACONH | 1.9 | 38 | 163h |
| U-$^2$H,$^{13}$C,$^{15}$N (refolded) | hCOCANH | 1.9 | 38 | 104h |
| U-$^2$H,$^{13}$C,$^{15}$N (refolded) | hcaCBcaCONH | 1.9 | 38 | 195h |
| U-$^2$H,$^{13}$C,$^{15}$N (refolded) | hNcocaNH | 1.9 | 38 | 99h |
| U-$^2$H,$^{13}$C,$^{15}$N (refolded) | hNCAcoNH | 1.9 | 38 | 193h |
| U-$^2$H,$^{15}$N, ketobutyrate, acetolactate | hCccNH TOCSY | 1.3 | 55.555 | 124h |
| U-$^2$H,$^{15}$N, ketobutyrate, acetolactate | hCCH TOCSY | 1.3 | 55.555 | 163h |
| U-$^{13}$C,$^{15}$N | CONCACX | 3.2 | 15 | 138h |
| U-$^{13}$C,$^{15}$N | CANCOCX | 3.2 | 15 | 112h |

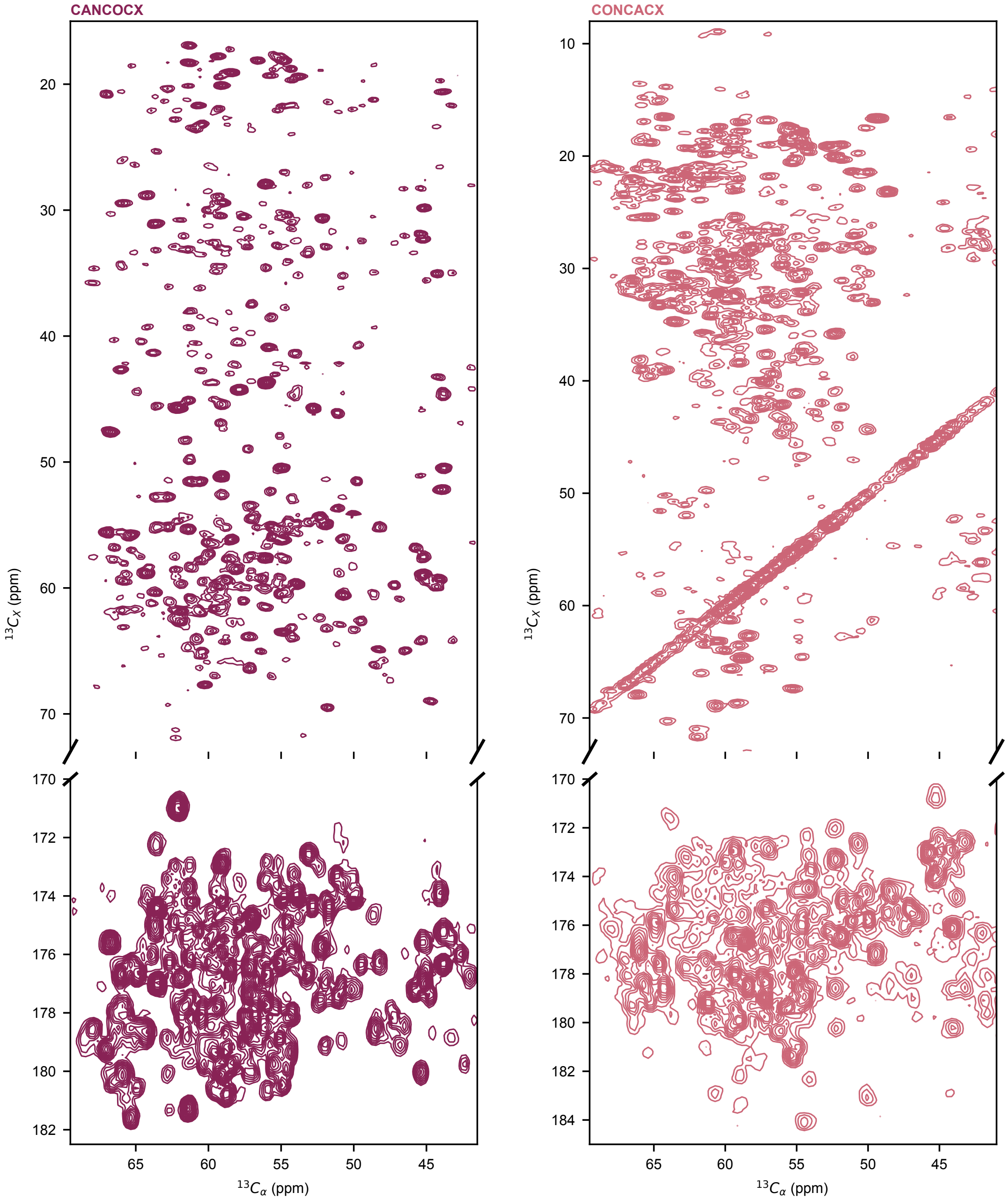

Figure S1: **$^{13}$C detected 4D MAS NMR spectra of sedimented *Ii*MDH recorded with a CPMAS CryoProbe.** Left: CANCOCX; right: CONCACX. Shown are projections along the dimensions N and CO.

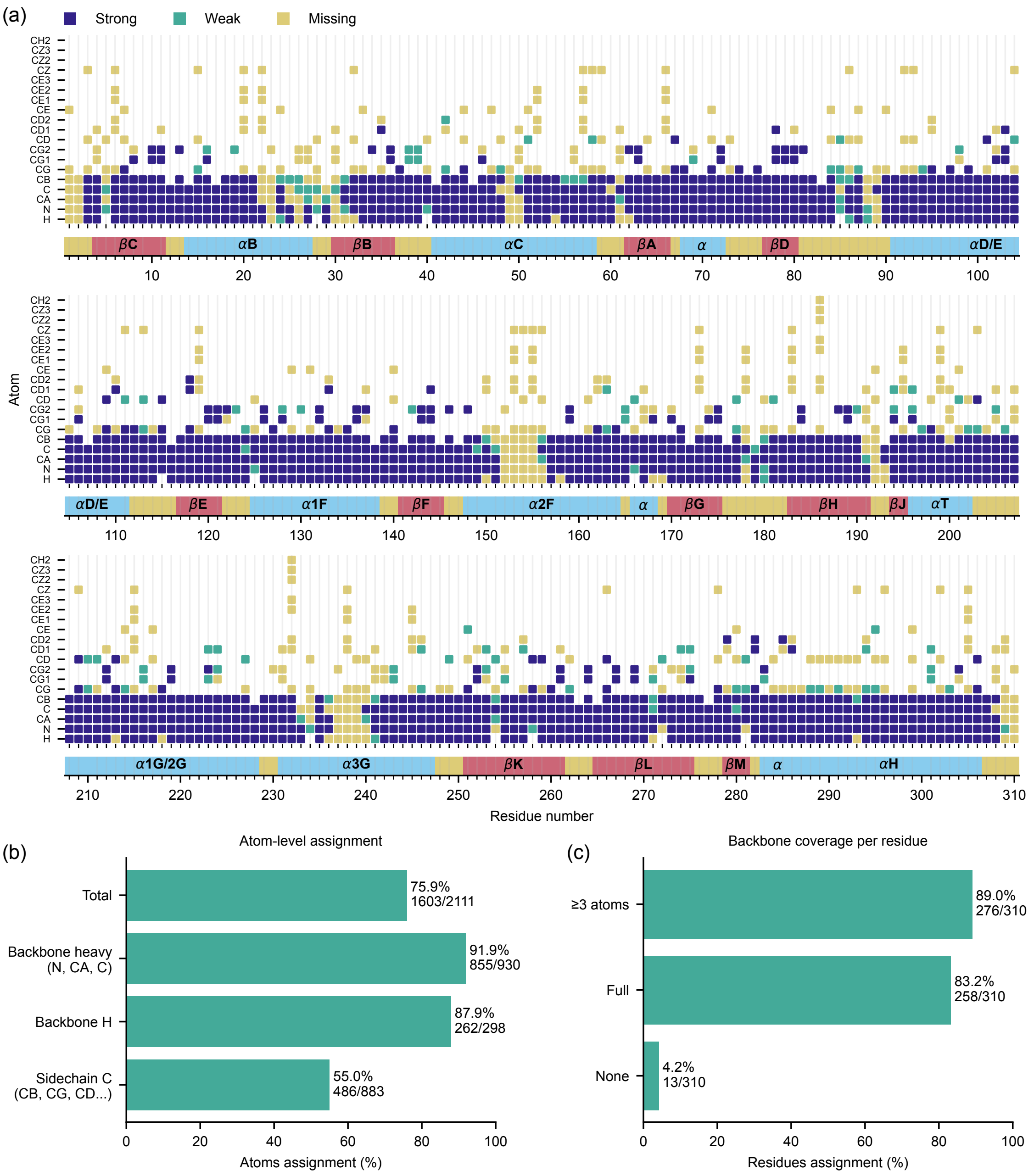

Figure S2: **MAS NMR assignment of backbone and sidechains.** (a) Extent of MAS NMR assignment. Weak assignments are supported by a single peak. (b) Assignment statistics, on a per-atom basis. (c) Assignment statistics, on a per-residue basis.

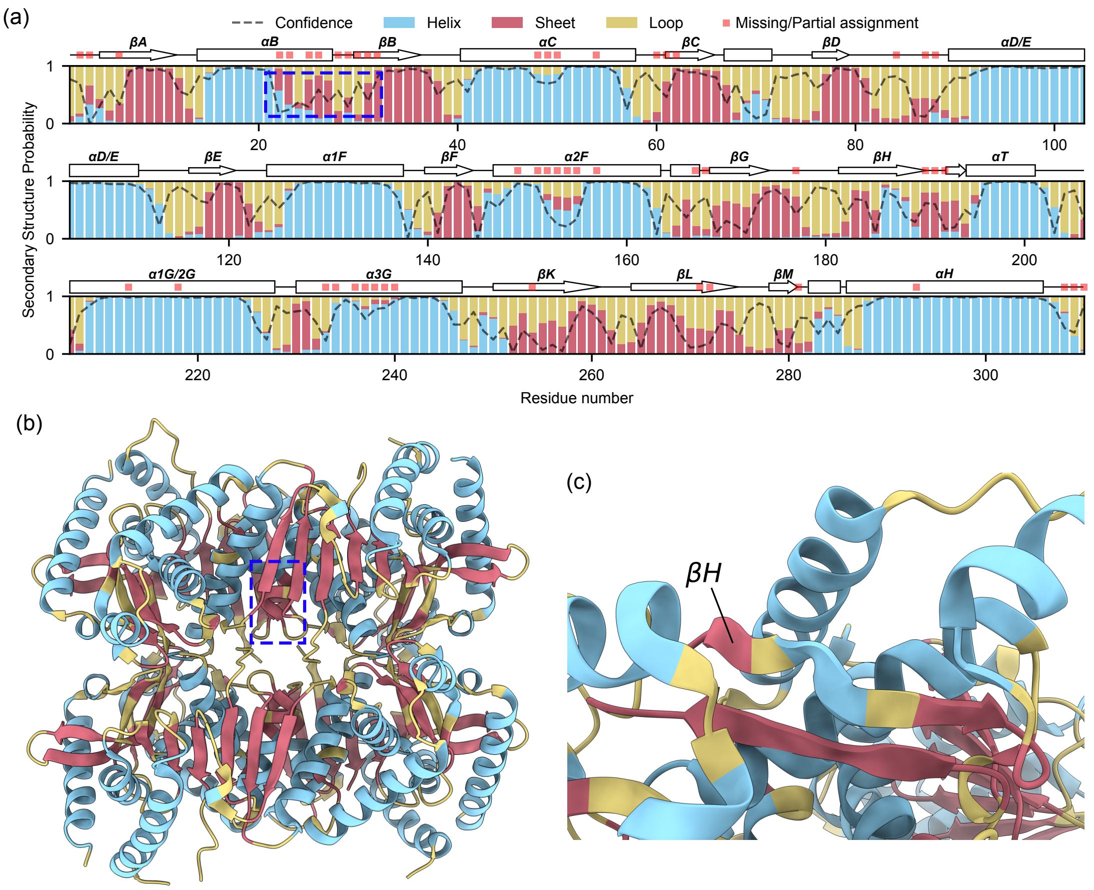

Figure S3: **Chemical shift-based prediction of secondary structures with TALOS-N[46].** (a) Secondary structure probability as predicted by TALOS-N using the MAS NMR assignments, with the prediction confidence. The secondary elements from the crystal structure (PDB: 6QSS) are reported over the stacked bar plot, as well as residues with missing/partial assignment of the backbone heavy-atoms. (b) *Ii*MDH crystal structure, colored based on the TALOS-N predictions, highlighting great agreement between the two, with the exception of helix αB (which is characterized by low confidence due to the lack of assignments). (c) Zoom on the βH strand. TALOS-N predicts a short turn in the middle of the strand, which is observed in the crystal structure too.

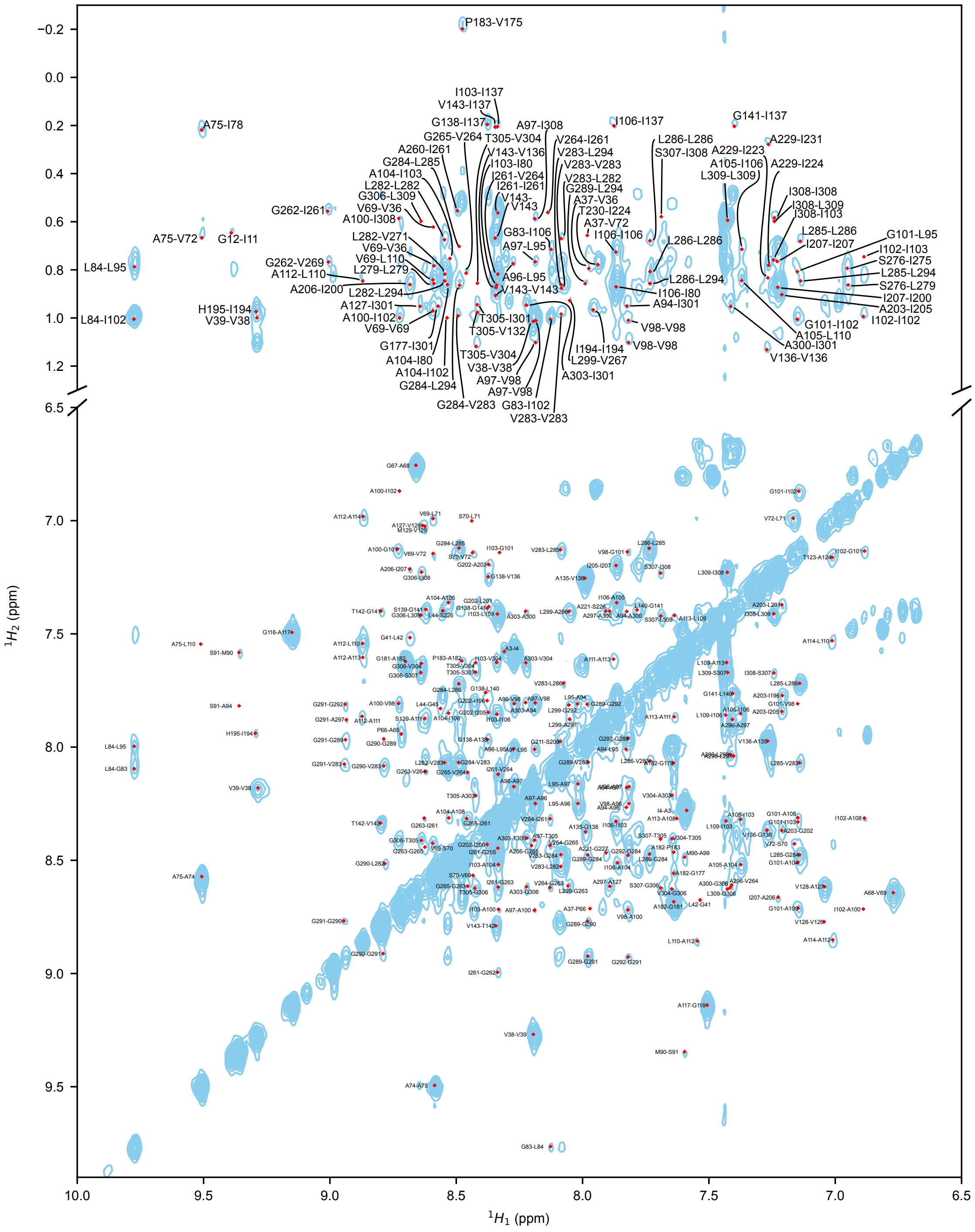

Figure S4: **H$^N$-(TROSY)-N-H NOESY spectrum projected along the N dimension**[49]. Through-space contacts between amide protons and amide/methyl protons can be detected in this spectrum.

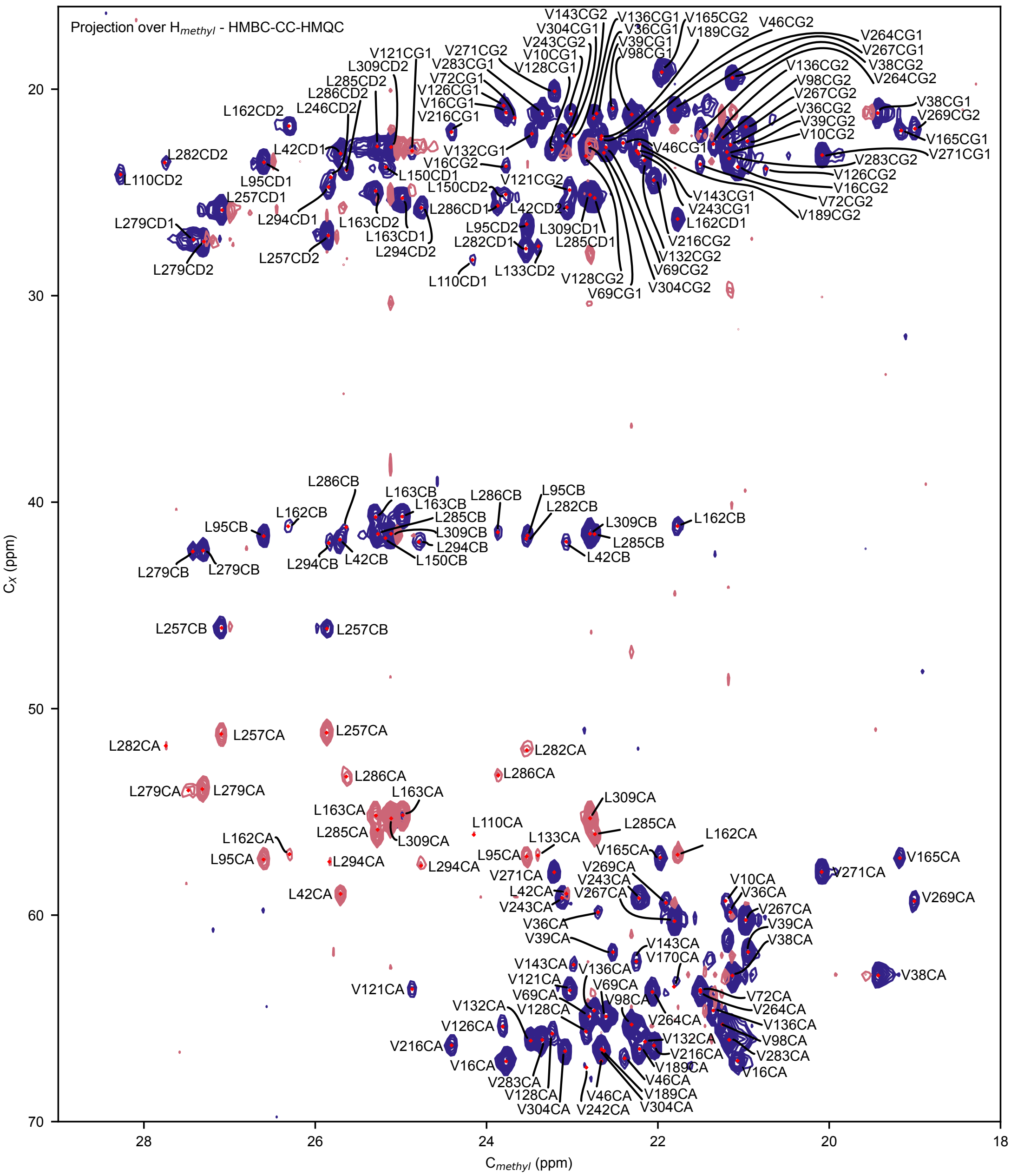

Figure S5: **HMBC-CC-HMQC spectrum projected along the H dimension.** In this spectrum, Leu-$\delta_1$/-$\delta_2$ and Val-$\gamma_1$/-$\gamma_2$ methyl group can be connected – through-bond – to their $C_\alpha/C_\beta$ and $C_\alpha$, respectively. The sample was produced using α-ketoisovalerate as a precursor[47].

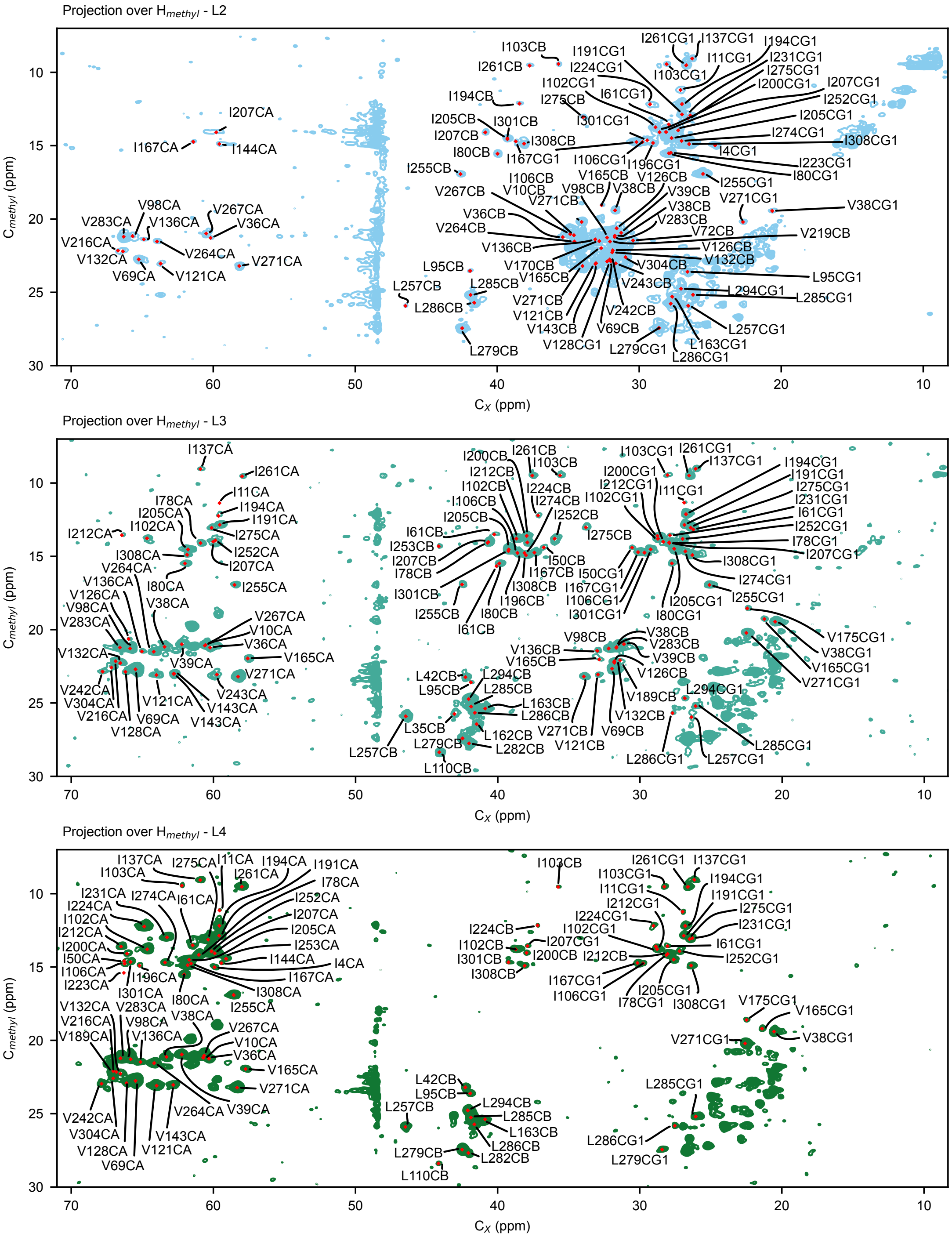

Figure S6: **TOCSY experiments projected along the H dimension.** We measured three TOCSY experiments correlating methyl groups with other carbon atoms[48]. Each experiment had a different mixing time (labeled L2, L3 and L4, corresponding to different numbers of FLOPSY cycles, ca. 11.7 ms, 17.7 ms and 23.5 ms). These resulted in different extents of transfer, favoring either the directly bonded (e.g., Ile-$\gamma_1$ in L2) or the further remote carbons (e.g., Ile-$\alpha$ in L4).

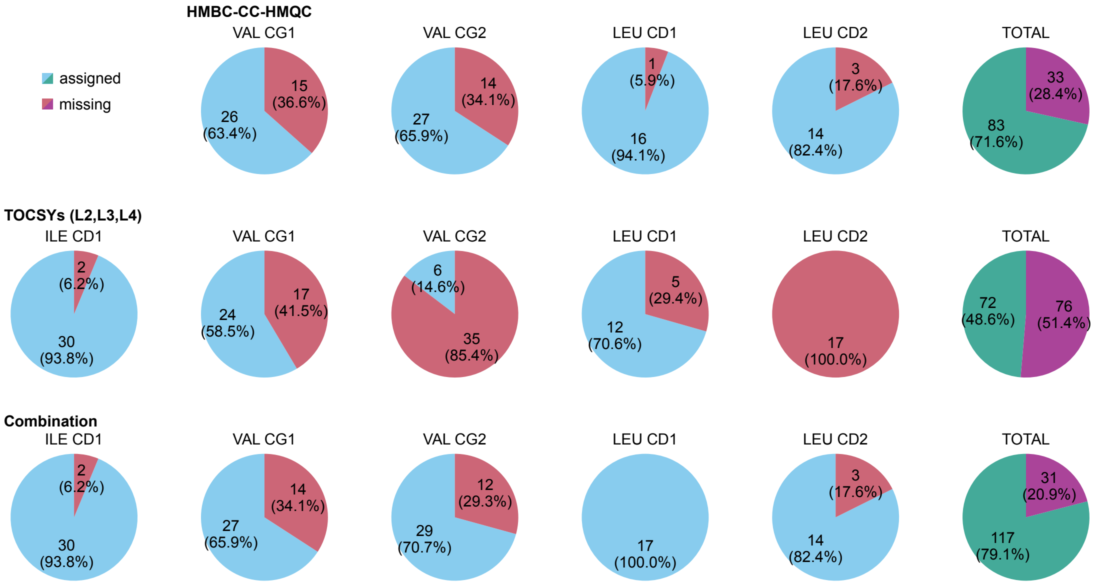

Figure S7: **Assignment of different methyl group classes in solution-NMR through-bond experiments.** The HMBC-CC-HMQC experiment recorded on the α-ketoisovalerate-labeled sample yielded good coverage of Val and Leu methyl groups. The TOCSYs experiments for the ketobutyrate/acetolactate-labeled sample performs worse (expecially for Val γ2 and Leu δ2), but nicely complements the previous approach with the Ile δ1 assignments (which are not accessible with the former labeling scheme).

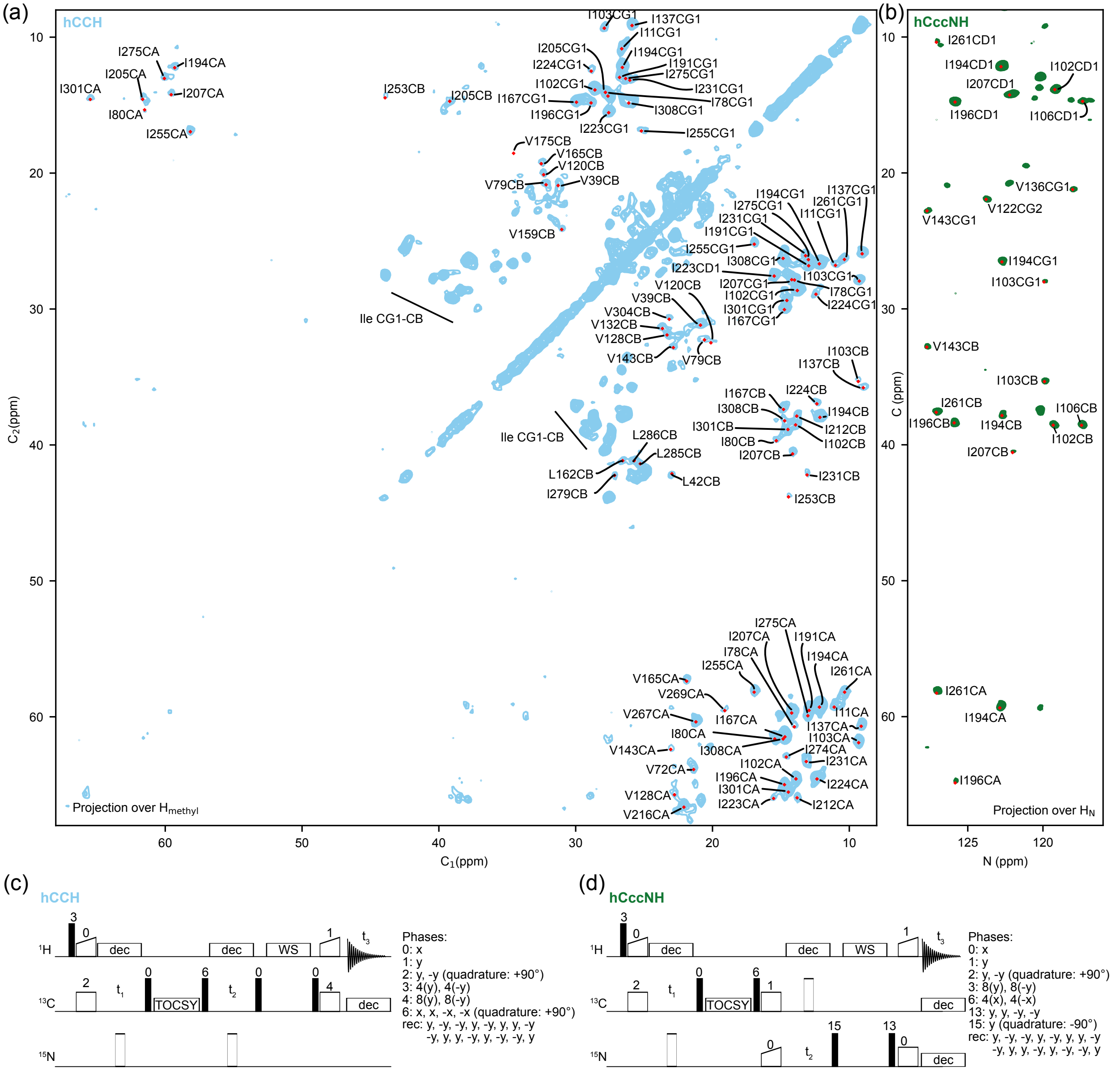

Figure S8: **TOCSY experiments for methyl assignment in the solid state**, applied to a sample with linear $^{13}$C chains in Ile, Leu and Val; the sample was the same as the one for the TOCSY experiment in solution shown in Fig. S6, but sedimented into a 1.3 mm rotor. (a) Projection of a 3D hCCH experiment along the methyl $^{1}$H dimension. The peak labels show the identity of the non-methyl $^{13}$C atom; the atom label not explicitly shown is Cδ1 (Ile, Leu) or Cγ1 (Val). (b) Projection of the hCccNH experiment, which connects the methyl carbon to the amide nitrogen and hydrogen frequencies. (c) Pulse sequence for the hCCH experiment, akin to previous implementations of HCC correlation experiments in solution[73] and in solids[74,75], uses cross-polarization steps (indicated with a ramp on the $^{1}$H channel) for transfer from $^{1}$H to $^{13}$C and back (3 ms, RF fields of ca. 40 kHz on $^{13}$C and 92 kHz (midpoint) on $^{1}$H with a 90-100 linear ramp), and a TOCSY mixing scheme for $^{13}$C-$^{13}$C transfer (DIPSI3[76], 21.7 ms, 10 kHz). (d) Pulse sequence for the hCccNH experiment, using cross-polarization steps ($^{1}$H-$^{13}$C: 3 ms, 40 kHz on $^{13}$C, 92 kHz (midpoint) on $^{1}$H with a 90-100 linear ramp; $^{13}$C-$^{15}$N: 9 ms, 13.7 kHz on $^{13}$C, 40 kHz (midpoint) on $^{15}$N with a linear 90-100 ramp; $^{15}$N-$^{1}$H: 1 ms, 91.2 kHz on $^{1}$H with a linear 100-90 ramp, 39 kHz on $^{15}$N), and a DIPSI-3 (21.7 ms, 10 kHz) $^{13}$C-$^{13}$C mixing as above. Pulse sequences and parameter files are available free of charge as supplementary material with this publication.

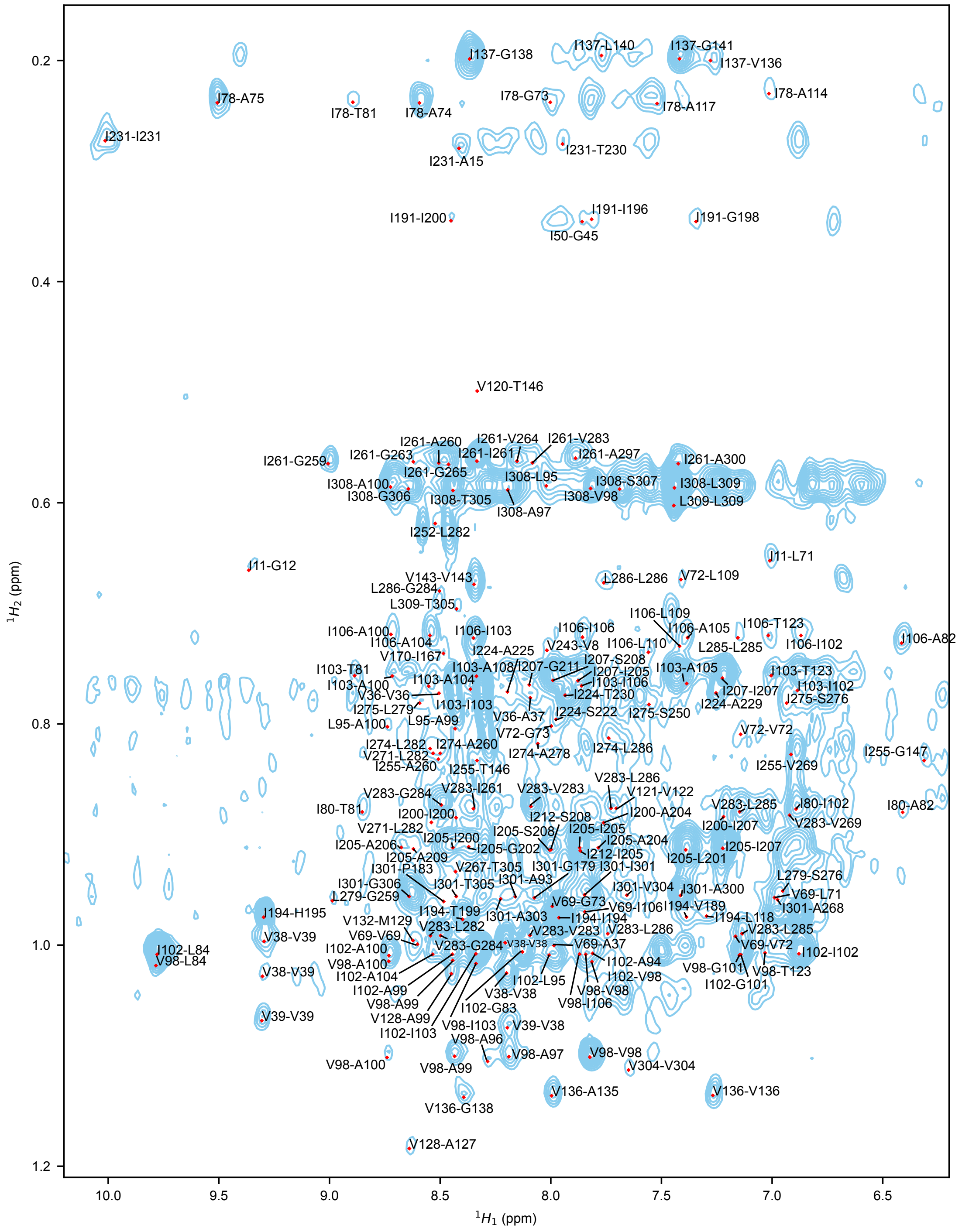

Figure S9: **H$^{C}$-C-H NOESY spectrum projected along the C dimension.** Through-space contacts between methyl and amide protons can be detected in this spectrum.

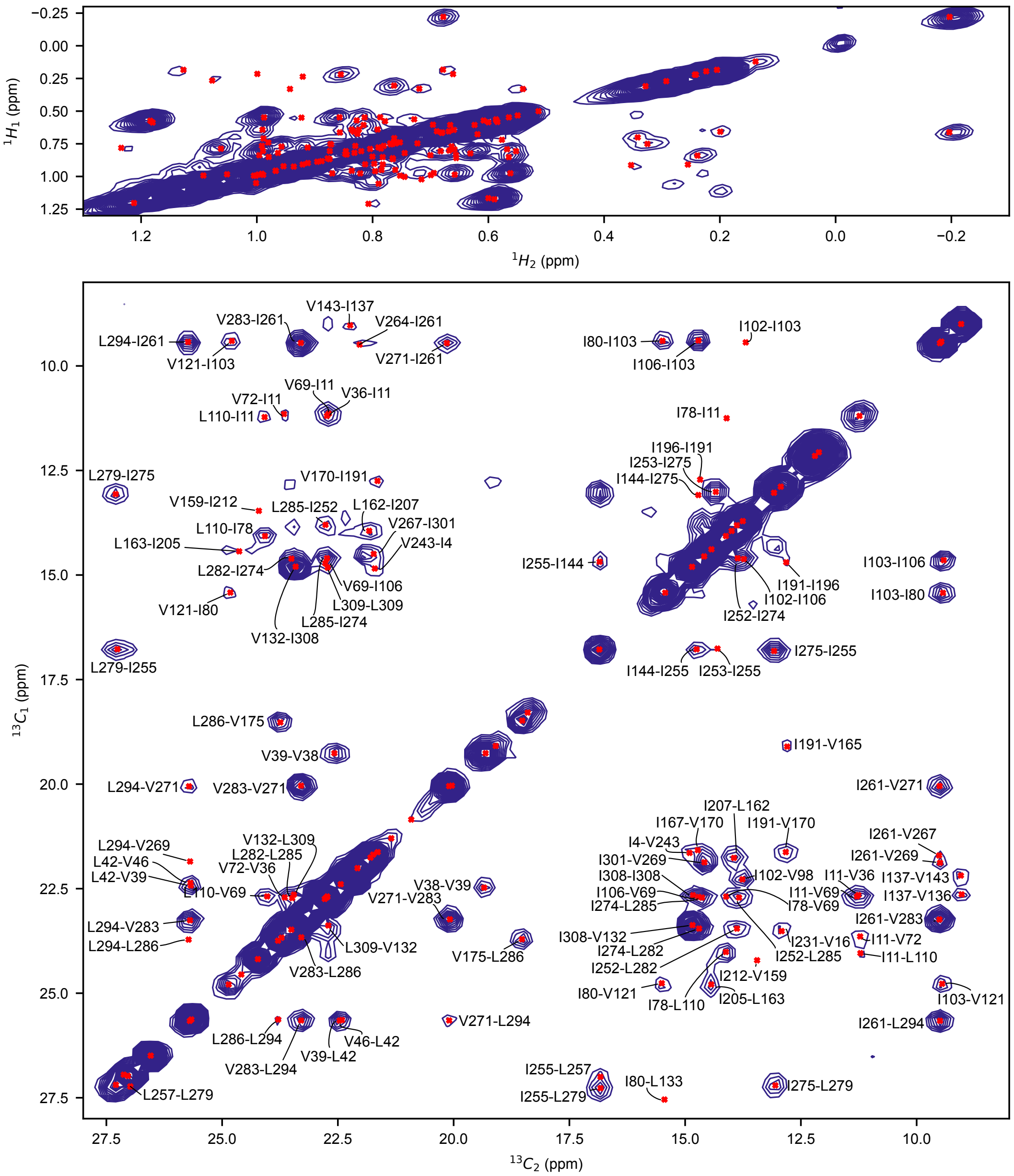

Figure S10: **H-H-C-C NOESY spectrum reporting on through-space Ile, Leu and Val methyl contacts.** Top: H-H plane (projection along the two C axes). Bottom: C-C plane (projection along two H axes).

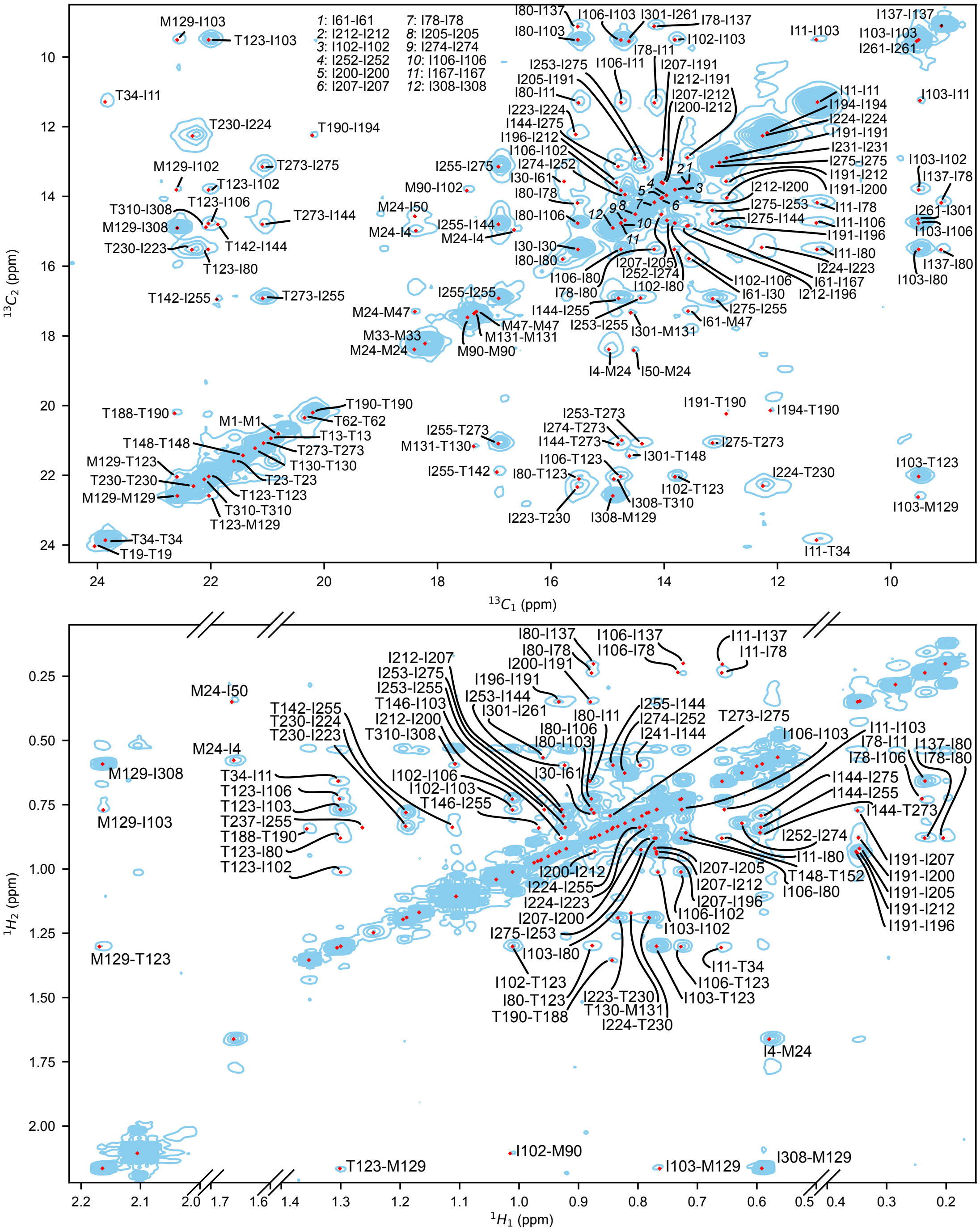

Figure S11: **NOESY spectra reporting on Ile, Met and Thr methyl contacts.** Top: H-C-C (projected along the H dimension). Bottom: H-H-C (projected along the C dimension).

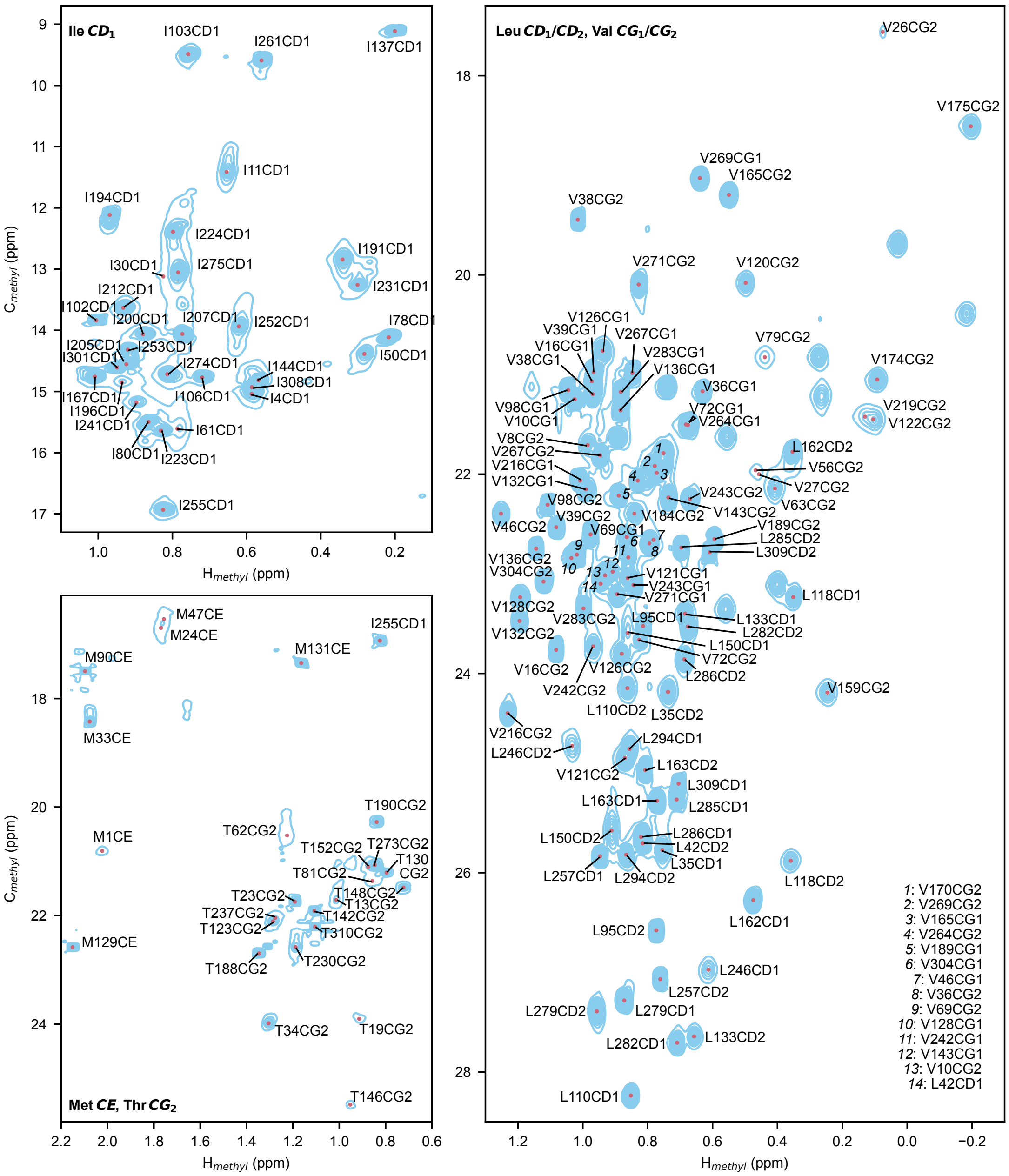

Figure S12: **Two-dimensional CH spectra of methyl groups.** Top-left: Ile-$\delta_1$. Bottom-left: Met-$\varepsilon$ and Thr-$\gamma$. Right: Leu-$\delta_1$/-$\delta_2$ and Val-$\gamma_1$/-$\gamma_2$ for mutant V168A.

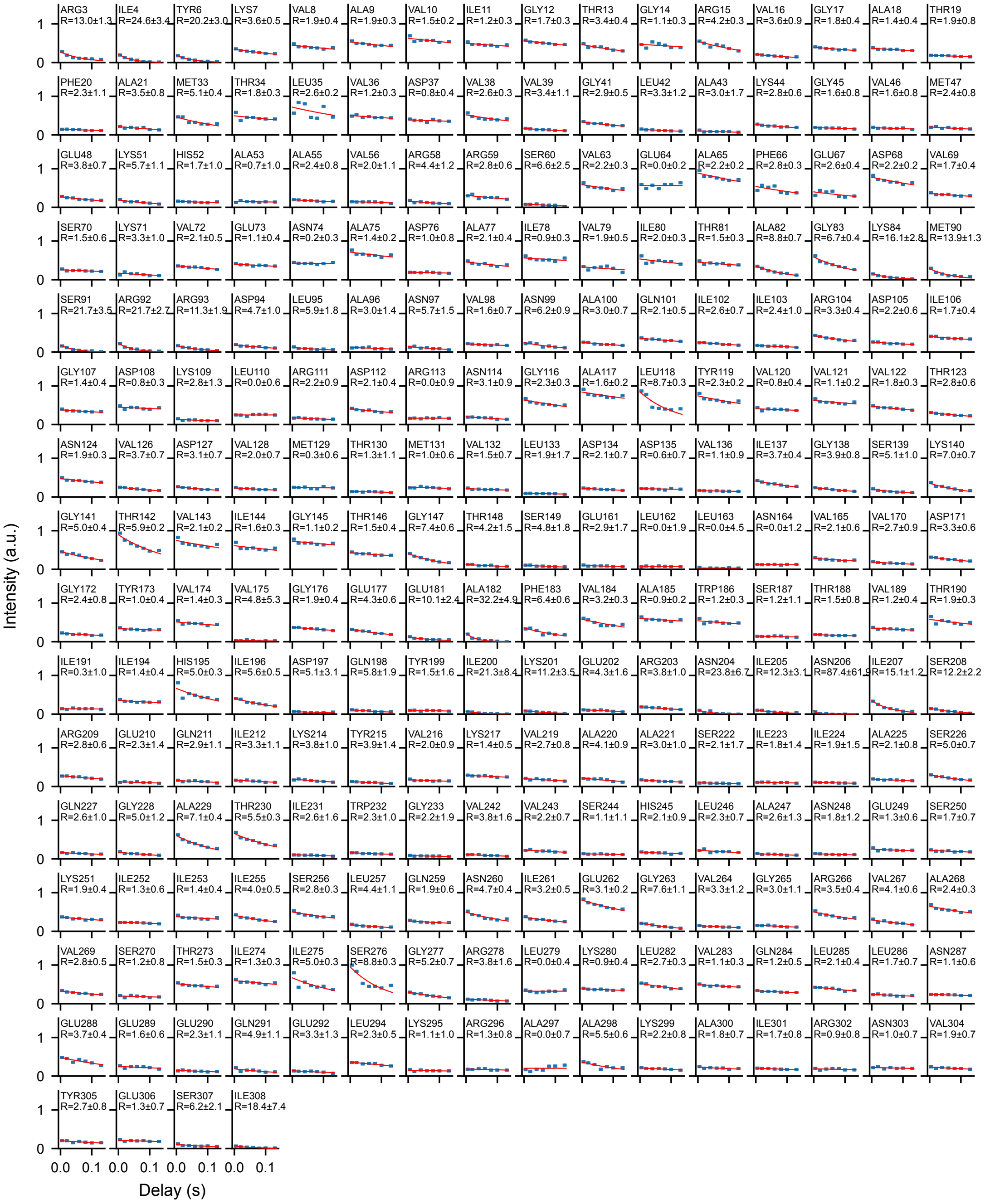

Figure S13: **MAS NMR $R_{1\rho}$ relaxation decays reporting on µs-ms dynamics.** The spinlock was set to 10 kHz; relaxation delays were 5, 20, 40, 60, 80, 100 and 130 ms.

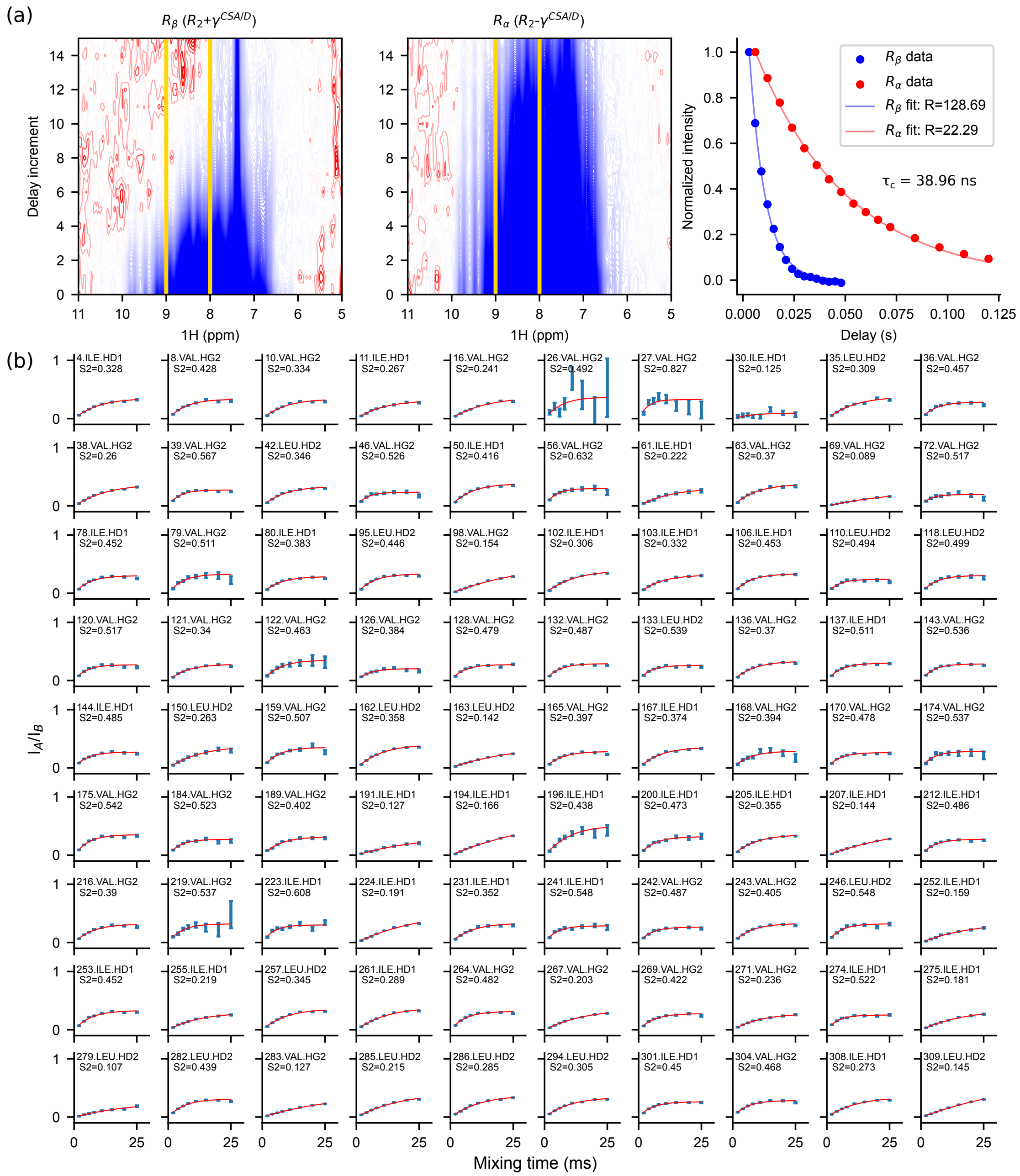

Figure S14: **$^{15}$N-TRACT and 3Q-relaxation buildups.** (a) $R_{2,\beta}$ and $R_{2,\alpha}$ decays with mono-exponential fits. $\tau_c$ (38.96 ns) was determined as in equation (6). (b) 3Q-relaxation buildups for Ile, Leu and Val residues. $S^2$ was determined as in equations (1-3). Relaxation delays were 2, 4, 6, 8, 11, 15, 20 and 25 ms.

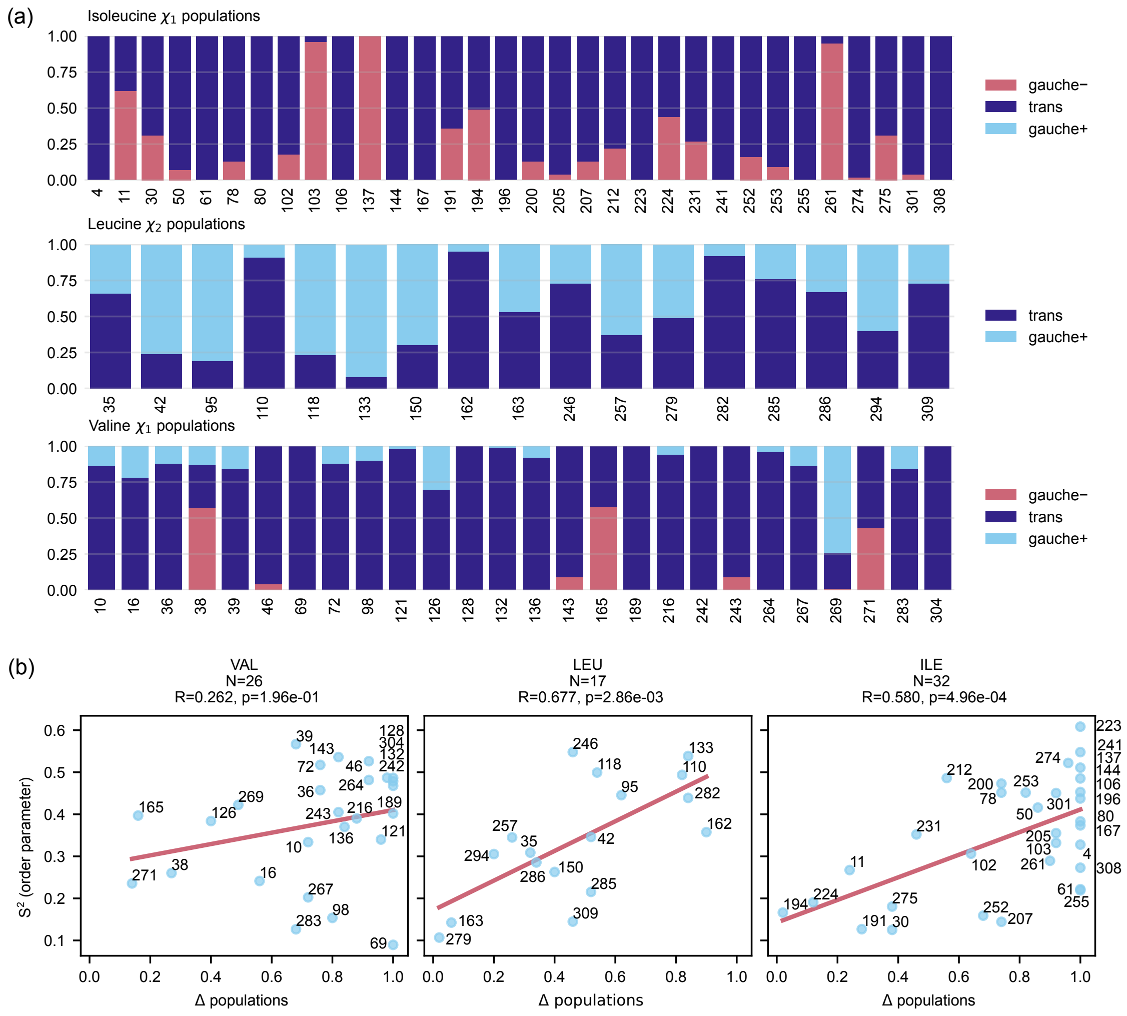

Figure S15: **Chemical shift-based prediction of rotamer populations.** (a) Isoleucine $\chi_1$, Leucine $\chi_2$ and Valine $\chi_1$ populations, predicted based on the $^{13}$C chemical shift of methyl groups[60–63]. (b) Correlation between methyl groups order parameters determined by 3Q-relaxation and variance in rotamer populations.

(a)

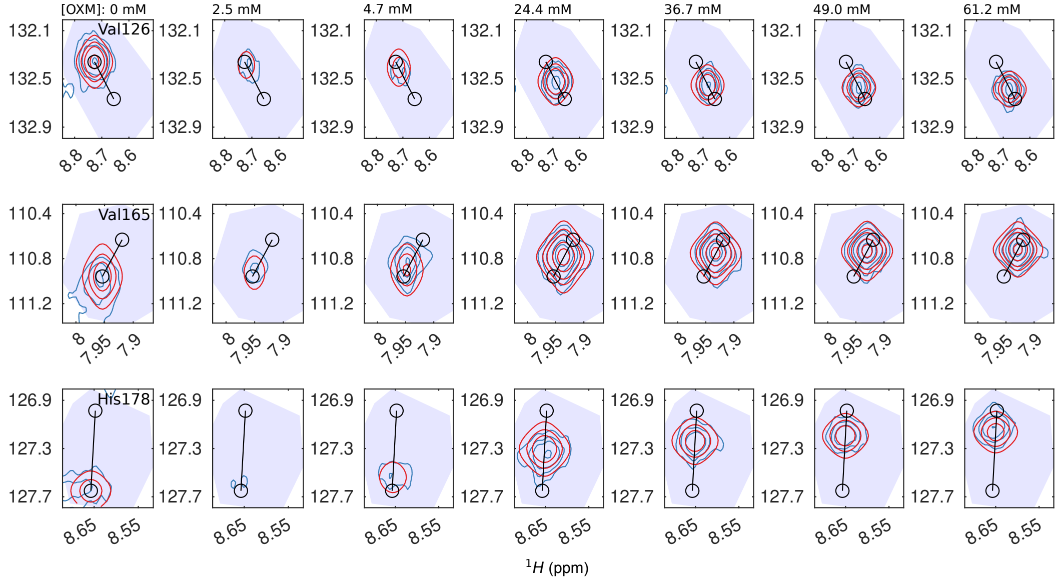

(b)

**Two-state ligand binding**

$P + L \rightleftharpoons PL$

$P_0$: protein concentration (e.g. $\mu$M)
$L_0$: ligand concentration (e.g. $\mu$M)

$K_d$: dissociation constant (e.g. $\mu$M)
$k_{\text{off}}$: dissociation rate ($s^{-1}$)

(c)

| PARAMETER NAME | FITTED VALUE | STD ERROR |
|---|---|---|
| Kd | 22484.265 | 1432.604 |
| koff | 999.080 | 173.500 |

Figure S16: **Two-dimensional lineshape analysis for $K_d$ estimation with NMR-TITAN[66].** (a) Raw spectra and NMR-TITAN fits for Val126, Val165 and His178, throughout the oxamate titration. (b) A two-state ligand binding model was chosen to fit the dissociation constant ($K_d$, in µM) and rate ($k_{off}$, in $s^{-1}$). (c) Fitted parameters and errors (bootstrapping method, 100 replicas).
